# The Importance of Mesozooplankton Diel Vertical Migration for Sustaining a Mesopelagic Food Web

**DOI:** 10.1101/642975

**Authors:** Thomas B. Kelly, Peter C. Davison, Ralf Goericke, Michael R. Landry, Mark D. Ohman, Michael R. Stukel

**Author notes:** Correspondence to Thomas B. Kelly.

## Abstract

We used extensive ecological and biogeochemical measurements obtained from quasi-Lagrangian experiments during two California Current Ecosystem Long-Term Ecosystem Research cruises to analyze carbon fluxes between the epipelagic and mesopelagic zones using a linear inverse ecosystem model (LIEM). Measurement constraints on the model include ^14^C primary productivity, dilution-based microzooplankton grazing rates, gut pigment-based mesozooplankton grazing rates (on multiple zooplankton size classes), ^234^Th:^238^U disequilibrium and sediment trap measured carbon export, and metabolic requirements of micronekton, zooplankton, and bacteria. A likelihood approach (Markov Chain Monte Carlo) was used to estimate the resulting flow uncertainties from a sample of potential flux networks. Results highlight the importance of mesozooplankton active transport (i.e., diel vertical migration) for supplying the carbon demand of mesopelagic organisms and sequestering carbon dioxide from the atmosphere. In nine water parcels ranging from a coastal bloom to offshore oligotrophic conditions, mesozooplankton active transport accounted for 18% - 84% (median: 42%) of the total carbon supply to the mesopelagic, with gravitational settling of POC (12% - 55%; median: 37%) and subduction (2% - 32%; median: 14%) providing the majority of the remainder. Vertically migrating zooplankton contributed to downward carbon flux through respiration and excretion at depth and via consumption loses to predatory zooplankton and mesopelagic fish (e.g. myctophids and gonostomatids). Sensitivity analyses showed that the results of the LIEM were robust to changes in nekton metabolic demands, rates of bacterial production, and mesozooplankton gross growth efficiency. This analysis suggests that prior estimates of zooplankton active transport based on conservative estimates of standard (rather than active) metabolism should be revisited.

**Contribution to the Field:** Understanding the flows of carbon within the ocean is important for predicting how global climate will shift; yet even after decades of research, the magnitude with which the ocean sequesters carbon is highly uncertain. One reason behind this uncertainty is that a variety of mechanisms control the balance between carbon input and carbon output within the ocean. The topic of this work is to inspect the role of biological organisms in physically transferring organic carbon from the surface to the deep ocean. As opposed to other mechanisms—such as sinking particles, the biological transfer of carbon is difficult to measure directly and is often quite variable, leading to large uncertainties. Here we use an extensive set of *in situ* observations off the coast of southern California to model the flow of carbon through the ecosystem. The model determined that in our study area nearly half of the total transfer of carbon from the surface ocean to deep was carried out by zooplankton that swim up to the surface each night to feed. This finding has direct implications for global carbon budgets, which often underestimate this transfer of carbon.

## 1. Introduction

Although mesopelagic food webs are believed to depend entirely on productivity generated in the euphotic zone, reconciling mesopelagic metabolic demand with estimates of export has been challenging (Burd et al., 2010; del Giorgio and Duarte, 2002; Hannides et al., 2015; Henson et al., 2011; Steinberg et al., 2008). Due to large uncertainties in rate measurements for meso- and bathypelagic organisms as well as low sampling resolution, steady-state budgets must either report wide ranges or otherwise exclude some processes, such as mortality and defecation of diel vertical migrators at depth. Even among recent studies, global carbon export budgets have been highly variable (Boyd and Trull, 2007; Henson et al., 2011, 2015; Laws et al., 2011; Siegel et al., 2014). Compounding this issue, several analyses have reported carbon demands by mesopelagic bacteria alone that exceed calculated carbon export (Burd et al., 2010; Ducklow and Harris, 1993), sometimes by an order of magnitude (Steinberg et al., 2008). This apparent imbalance between the carbon supply to the mesopelagic and the estimated metabolic demand suggests either export estimates fail to capture important dynamics or that metabolic calculations are grossly overestimated (Burd et al., 2010). Notably, the most prominent study to date that has purported to show a balanced epipelagic-mesopelagic carbon budget did so by assuming much lower bacterial carbon:leucine ratios than used in other estimates of bacterial production (Giering et al., 2014). Hence, their results may have been derived from misinterpretation of bacterial leucine uptake.

Recent work has demonstrated that diel vertical migrators are important for net transfer of organic carbon from the euphotic zone to the mesopelagic, a transfer not measured with traditional carbon export methods (Morales, 1999; Steinberg et al., 2000). Since export by mesozooplankton is not captured by sediment traps or chemical disequilibria, we must rely on logistically difficult biomass-based estimates or on indirect modeling syntheses. For example, using remote sensing fields and a size-structured ecosystem model, Archibald et al. (2019) found that global zooplankton diel vertical migration (DVM) can increase export production by 14% annually. This is consistent with previous modeling exercises based on zooplankton behavior (Bianchi et al., 2013) and community size structure (Aumont et al., 2018). Zooplankton behavior models argue that for DVM to be evolutionarily advantageous (Cohen and Forward, 2009), the energy expenditure should be offset by a commensurate reduction in predation risk. Using this modeled-behavior approach, Hansen and Visser (2016) found that 16% - 30% mid-latitude export production in the North Atlantic was due to DVM mesozooplankton. Each of these models note sensitivities to zooplankton biomass and the fraction of the zooplankton population that undergoes DVM, which are ecosystem metrics that are difficult to generalize.

Linear inverse ecosystem models (LIEM) have been shown to be a versatile and robust framework for integrating a wide range of ecosystem data (Gontikaki et al., 2011; Sailley et al., 2013; Stukel et al., 2018b; van Oevelen et al., 2012; Vézina et al., 1988). An LIEM combines an ecosystem network with observations and generalized constraints to determine possible energy flows through the ecosystem. Unlike a forward model (e.g., an NPZ model; Franks, 2002), the relationships between organisms are not prescribed by functional responses of model state variables (e.g., assuming a Monod functional form controls phytoplankton nutrient uptake responses or an Ivlev grazing formulation). Instead, the model includes all possible combinations of fluxes that are compatible with the input constraints, and the most likely ecosystem structure is then retrieved based on a random walk through the solution space (van den Meersche et al., 2009). This inverted approach has the advantage of not requiring *a priori* assumptions of ecological relationships but instead relies on many independent constraints on the food web.

The California Current Ecosystem (CCE) is an eastern boundary current upwelling biome with extensive temporal and spatial variability. As a result of high mesozooplankton biomass and strong DVM (Ohman and Romagnan, 2016; Powell and Ohman, 2015; Stukel et al., 2013), we expect a substantial contribution to export production by diel vertical migrators and a commensurately important role in satisfying the carbon demand of the mesopelagic. Stukel et al. (2013) suggested that active transport could be responsible for 1.8 – 29% of total export in the CCE. However, their study focused only on active transport fluxes due to zooplankton respiration and used conservative assumptions. To more thoroughly investigate the potential importance of active transport, we designed a two-layer LIEM, which includes non-living organic matter, primary producers, zooplankton and planktivorous nekton organized into two layers: an epipelagic and a mesopelagic ecosystem. Using extensive data from two cruises of the CCE Long-Term Ecological Research (LTER) Program in the southern California Current region, our LIEM data synthesis suggests that active transport of carbon from the epipelagic down to depth is a significant mechanism supporting the mesopelagic carbon demand. Although previous studies have indicated that active transport may be responsible for 10% - 30% of total carbon flux (Archibald et al., 2019; Aumont et al., 2018; Bianchi et al., 2013; Hansen and Visser, 2016; Yebra et al., 2005), our LIEM suggests that 20% - 80% of carbon export in the CCE may be the result of DVM by mesozooplankton.

## 2. Materials and methods

### 2.1. Ecosystem data

The data presented here (Appendix A) were collected during two cruises of the California Current Ecosystem Long Term Ecological Research (CCE LTER) program (P0704 in April 2007; P0810 in Oct. 2008). On these cruises, *in situ* drift arrays were used for quasi-Lagrangian tracking of water parcels for periods of 3-5 days (Landry et al., 2009, 2012), while the water column was repeatedly sampled for the following variables: CTD-derived physical data, phytoplankton diversity and biomass (flow cytometry, epifluorescence microscopy, and pigment analyses, (Taylor et al., 2012)), primary production (H^14^CO_3_-uptake, (Morrow et al., 2018)), mesozooplankton biomass and community analyses (paired day-night bongo and Multiple Opening and Closing Net with Environmental Sampling System, MOCNESS net tows, (Ohman et al., 2012; Powell and Ohman, 2012), microzooplankton biomass (epifluorescence microscopy), microzooplankton grazing (dilution method, Landry et al., 2009), mesozooplankton grazing (gut pigment methods, Landry et al., 2009), meso- and epipelagic micronekton biomass and metabolic demands (see Section 2.1.1; Oozeki net trawls, multi-frequency EK60 echosounder, and individual-based metabolic model (Davison et al., 2013, 2015), bacterial production (^3^H-leucine uptake, Samo et al., 2012), and gravitational particle export (sediment traps and ^234^Th:^238^U disequilibrium, Stukel et al., 2013). The use of a quasi-Lagrangian sampling framework also allowed us to assess net rates of change of phytoplankton biomass. Bulk rates and associated errors for the 3-5 day cycles were calculated by averaging vertically-integrated rates or biomasses for each experimental cycle. The data and detailed methods can be found on the CCE LTER Datazoo website (http://oceaninformatics.ucsd.edu/datazoo/data/ccelter/datasets) and/or in published manuscripts cited above.

The quasi-Lagrangian experiments (hereafter ‘cycles’) spanned much of the physical, chemical, and ecological variability of the CCE domain (Table 1, Fig. 1) which allowed us to classify cycles according to nutrient conditions, the primary driver of ecosystem variability within the CCE (Landry et al., 2012). Cycle classification was defined as: nutrient-limited cycles which were conducted in off-shore, low nutrient regions (P0704-2, P0810-2, P0810-6); transition region cycles which were characterized by low surface nutrient concentrations and intermediate NPP and biomass (P0810-1, P0810-3, P0810-4); and upwelling cycles in which surface nutrient concentrations and phytoplankton growth rates were highest (P0704-1, P0704-4, P0810-5; Table 1).

**Table 1.** Overview of conditions for each cycle along with the attributed classifications: upwelling, transition region, and nutrient limited.

| Cycle | Classification | Surface Chl<br>(μg Chl a L<sup>-1</sup>) | <sup>14</sup>C Primary<br>Productivity<br>(mg C m<sup>-2</sup> d<sup>-1</sup>) | Mesozooplankton<br>Biomass<br>(mg C m<sup>-2</sup>) |
| --- | --- | --- | --- | --- |
| P0704-1 | *Upwelling* | 1.35 | 1,233 | 2,695 |
| P0704-2 | *Nutrient Limited* | 0.22 | 587 | 391 |
| P0704-4 | *Upwelling* | 0.99 | 2,314 | 1,715 |
| P0810-1 | *Transition region* | 0.45 | 554 | 740 |
| P0810-2 | *Nutrient Limited* | 0.20 | 484 | 528 |
| P0810-3 | *Transition region* | 0.72 | 892 | 923 |
| P0810-4 | *Transition region* | 1.05 | 674 | 832 |
| P0810-5 | *Upwelling* | 1.47 | 1,672 | 1,098 |
| P0810-6 | *Nutrient Limited* | 0.22 | 325 | 628 |

**Figure 1.**
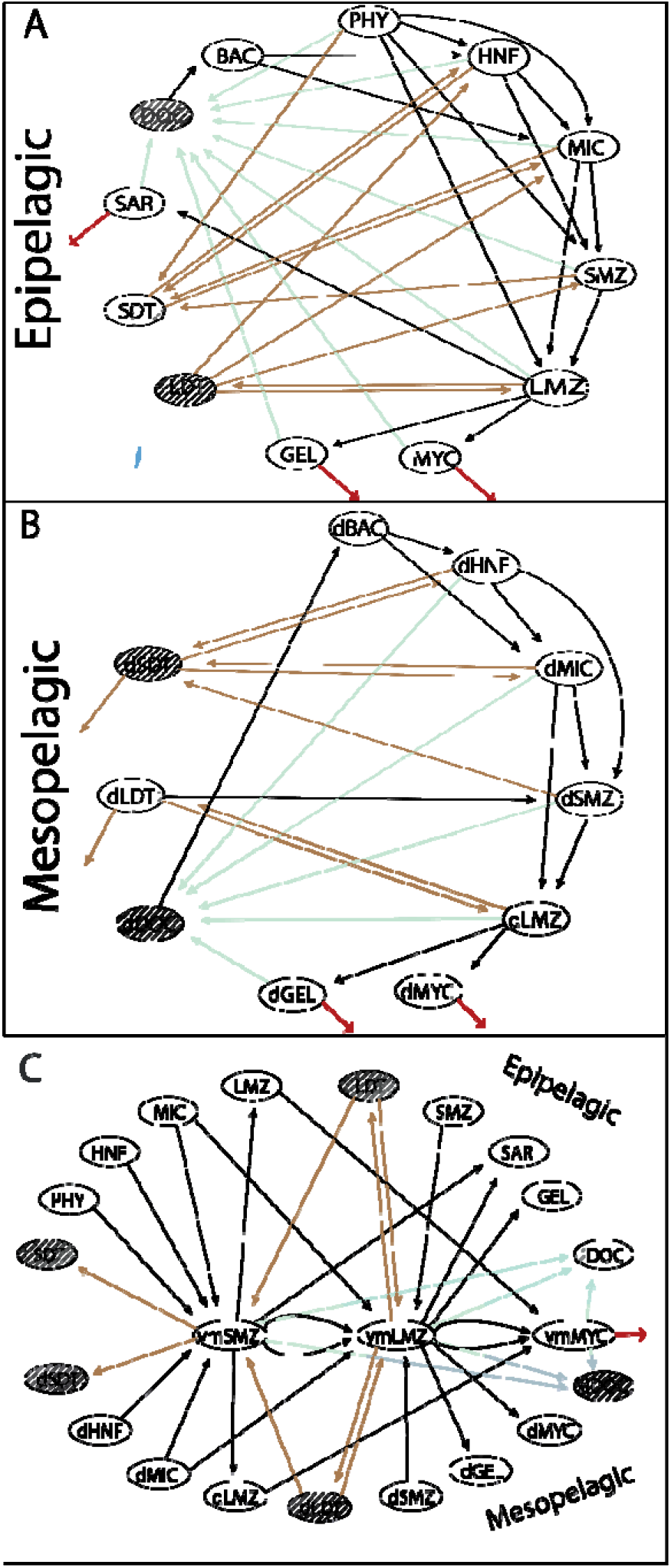
Schematic of model structure organized into distinct layers where arrows indicate a model flow. (A) Epipelagic organisms and POC. Arrows are colored for convenience: direct flows to the mesopelagic = blue, flows to HTL = red, flows to and from detritus = brown, and flows to DOC or dDOC = green. Shaded compartments indicate no losses to respiration. (B) Same as (A) for the mesopelagic. (C) Showing the connectivity between the epipelagic and mesopelagic layers via vertical migrators. See Table 2 for abbreviations.

#### 2.1.1. Phytoplankton, Bacteria, and Protist Constraints

Daily *in situ* primary productivity measurements using H^14^CO_3_^−^ uptake (^14^CPP) were conducted at 6-8 depths spanning the euphotic zone using 4 L incubations subsampled in triplicate (Morrow et al., 2018). A 250 mL dark bottle was used to correct for non-photosynthetic ^14^C uptake. Contemporaneously, *in situ* dilution experiments, using the two-treatment approach of Landry et al. (2008), were conducted to measure protistan zooplankton grazing rates (Landry et al., 2009). Euphotic zone primary production and protistan zooplankton gazing rates were vertically integrated and averaged by cycle.

**Table 2.**
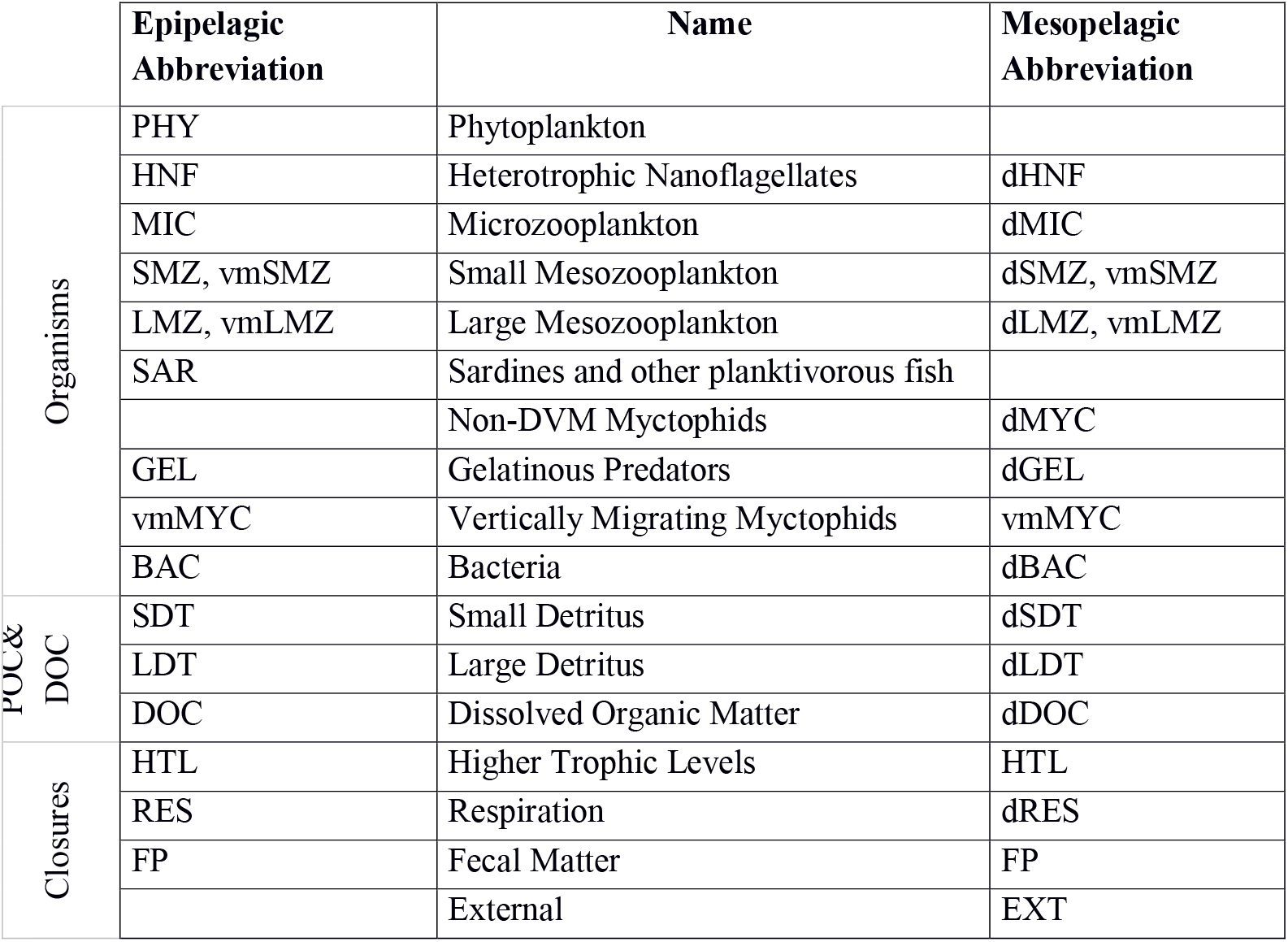
Names and abbreviations of all model compartments. An abbreviation in the left column indicates inclusion in the epipelagic, while an abbreviation in the right column indicates inclusion in the mesopelagic. Each abbreviation is a distinct compartment in the LIEM with the prefix ‘vm’ signifying vertical migration and ‘d’ signifying the mesopelagic.

Rates of ^3^H-leucine incorporation into bacteria were measured in triplicate at multiple depths during each cycle (Samo et al., 2012). Each profile was vertically integrated and then averaged by cycle in order to determine epipelagic bacterial production rates. Additionally, an upper and lower bound for mesopelagic bacterial production was calculated by integrating a bacterial production attenuation curve and then scaling by the epipelagic bacterial production (Eq. 1).

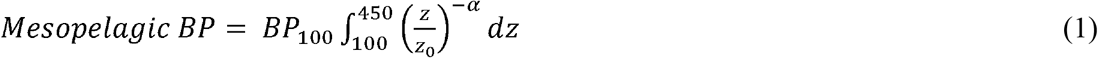

where *BP*_*100*_ is the measured BP rate at 100 m and α (BP attenuation factor) = 1.47 (Yokokawa et al., 2013) for the lower limit and α = 0 (i.e., no attenuation) for the upper limit.

#### 2.1.2. Mesozooplankton and Nekton Constraints

Data for the mesozooplankton constraints primarily comes from either day-night paired oblique bongo net tows through the epipelagic (for grazing rates) or day-night paired MOCNESS tows taken at 9 depth horizons spanning the upper 450 m (for biomass and metabolism estimates). MOCNESS samples were analyzed by ZooScan digital scanner (Gorsky et al., 2010; Ohman et al., 2012), vignettes provisionally classified using machine learning methods, then 100% manually validated. Organisms were sorted (Stukel et al., 2013) into groups including euphausiids, nauplii, copepods, appendicularians, siphonophores, and other crustaceans. For this study, we separated the mesozooplankton community into two size classes (<1 mm and >1 mm) of grazers and one compartment for gelatinous predators (siphonophores). We also partitioned the large and small mesozooplankton into non-vertically migrating epipelagic residents, vertical-migrators, or mesopelagic resident communities. Biomass of the non-vertically migrating, epipelagic mesozooplankton was calculated from the upper 100 m net during daytime tows while the non-vertically migrating, mesopelagic biomass was calculated based on the nighttime mesopelagic (100 m – 450 m) net tows (Stukel et al., 2013). We note that epipelagic biomass estimates are likely conservative due to net avoidance. Biomass estimates for the DVM mesozooplankton were calculated by averaging the difference in the night and day epipelagic biomass estimates with the difference in the day and night mesopelagic biomass estimates. This approach was used in order to be the most consistent with both the epipelagic and mesopelagic biomass estimates for non-vertically migrating biomass. For a list of abbreviations used for all model compartments, see Table 2.

Minimum respiration for each mesozooplankton group was calculated using published temperature-length-basal respiration relationships (Ikeda et al., 2001). Oxygen consumption was converted to carbon units using the scale factor 9.88 × 10^−3^ mg C d^−1^ (μL O_2_ hr ^−1^)^−1^. Mesozooplankton grazing on phytoplankton was calculated from gut pigment contents of oblique bongo net tow tows (202 µm mesh, D = 0.71m) and estimated gut passage rates (Dam and Peterson, 1988). Carbon-based grazing rates were then calculated from chlorophyll (Chl) consumption, and C:Chl ratios computed as the ratio of NPP to chlorophyll-specific growth rates obtained from the dilution experiments. Mesozooplankton grazing rates were size fractionated as above. Mesozooplankton gut contents samples were improperly frozen for P0810-5, P0810-6, and most of P0810-4. In order to provide estimates for these grazing rates, average grazing rates from the cycle with the same classification were used (e.g. P0810-5 was an upwelling cycle so grazing rates were averaged from the other upwelling cycles). Conservative uncertainty estimates were set to be 2x the error calculated by propagation of error. This higher level of uncertainty is a reasonable compromise given the data limitations. For additional details on gut pigment processing, see Landry et al. (2009).

Nekton biomass was estimated based on catches made by a 5 m^2^ Matsuda-Oozeki-Hu net trawl (Davison et al., 2013). For each station, epipelagic net tows were conducted at night after the ascent of the deep scattering layer. Preserved specimens from each net tow were identified to species and measured. Fish were classified as either non-vertical migrating or vertically migrating based on species. An individual based model was then used to determine metabolic rates and requirements for each nekton population: resident epipelagic, diel vertical migrant, and resident mesopelagic (Davison et al., 2013).

#### 2.1.3. Export Production

VERTEX-style sediment traps consisting of 8-12 tubes per depth were deployed and recovered at the start and end of each cycle (Knauer et al., 1979; Stukel et al., 2013). Tubes were filled with a hypersaline, poisoned brine solution. Upon recovery >200-µm swimming mesozooplankton taxa were manually removed during inspection under a stereomicroscope. Samples for C and N or C:^234^Th ratios were filtered through pre-combusted glass fiber and quartz filters, respectively, prior to analysis on a CHN elemental analyzer or a RISO beta multi-counter.

^234^Th:^238^U disequilibrium measurements were made at 12 depths spanning the upper 200 m at the start and end of each cycle using standard small-volume procedures (Benitez-Nelson et al., 2001; Pike et al., 2005). Thorium-234 export rates were then computed using a 1-box steady state model (Savoye et al., 2006). The C:^234^Th ratio measured from sediment trap particles was used to convert to carbon export. For additional details, see (Stukel et al., 2019).

Subduction of POC provides an alternative mechanism for the export of organic matter to the mesopelagic, that is not measured by either sediment traps or ^234^Th profiles, which only record gravitational settling of particles. A three-dimensional particle advection model was used to determine a range of possible subduction rates (Stukel et al., 2018c). The maximum and minimum estimates of particle subduction were used as bounds on two size-fractionated subduction flows within the LIEM.

### 2.2. Linear inverse model

We developed a LIEM for the CCE to investigate mechanisms of epipelagic-mesopelagic coupling. The LIEM consists of 140 flows (i.e., ecosystem fluxes, Supp. Table 2) and 24 compartments (i.e. standing stocks; Table 2) organized into two layers: the surface epipelagic and a deeper mesopelagic ecosystem (defined as 100 – 450 m depth to match with *in situ* measurements). The epipelagic and mesopelagic ecosystems consist of 73 flows and 64 flows, respectively, with four explicit flows (particle sinking and subduction) and three implicit flows (active transport) directly linking the two layers (Figure 1). Three vertically migrating compartments (small and large mesozooplankton and nekton) connect the epipelagic and mesopelagic through a transfer associated with DVM (i.e. respiration, excretion, and mortality). Constraints consist of 24 mass balance equations, 18 approximate equations (i.e. *in situ* rate measurements) and 133 inequalities, which are provided in the Online Supplement.

The 18 approximate equations are ecosystem observations, which can be directly compared to flows within the model (Appendix A). These equations are net primary productivity (NPP), phytoplankton biomass net rate of change, protistan grazing, size-fractionated grazing rates (<1-mm and >1-mm) for epipelagic resident and DVM mesozooplankton, sediment trap and ^234^Th-based export fluxes, bacterial production, and mesopelagic fish respiration, mortality and fecal pellet production rates. The model was provided an estimated value and associated uncertainty for each measurement.

Respiration, mesopelagic export, nekton fecal pellets, and losses to higher trophic levels were included as closure terms. Within the model every organism loses carbon to respiration, DOC excretion, and defecation or mortality to detritus/fecal pellets. Grazing was allowed between organisms whose ecological role and size-range permit grazing (e.g. small mesozooplankton graze on nano- and microplankton; sardines consume only >1-mm mesozooplankton). Mass balance was required for each compartment. All compartments were assumed to be at steady state except for PHY, for which changes in biomass were measured (via Chl-*a* proxy) during each cycle and incorporated into the model. This flexibility was essential to capture the bloom-phase of the ecosystem since dramatic shifts in Chl-*a* were observed during some cycles.

#### 2.2.1. Inequality constraints

The formulas used in the inequality constraints are provided in the Online Supplement. Upper and lower limit estimates of POC subduction from the epipelagic to the mesopelagic layer were taken from (Stukel et al., 2018c), and a minimum fecal pellet flux was assigned based on the assumption that recognizable fecal pellets in sediment trap material represented a lower limit on total fecal pellet flux. Minimum and maximum Gross Growth Efficiencies (GGE) and Absorption Efficiencies (AE) were assigned according to previously accepted literature values: 10% – 40% GGE for protistan zooplankton (HNF & MIC) and gelatinous predators (Straile, 1997); 10%-30% for mesozooplankton (Anderson et al., 2018); 5% – 30% for bacteria (del Giorgio and Cole, 1998); and 50% – 90% AE for all heterotrophs (Conover, 1966).

Minimum respiration requirements were considered as both active respiration and basal respiration. Active respiration was set as a fraction of ingestion, and basal respiration was set as a function of biomass and temperature. Valid solutions fulfilled both criteria. Diel vertical migrator biomass, as determined from MOCNESS net tows, was used to calculate a minimum respiration based on temperature. DOC excretion was required to be greater than 10% of ingestion (or 2% of NPP for phytoplankton) and less than respiration (or 35% of NPP). All inequality constraints are listed in Supplemental Table 1.

#### 2.2.2. Model solution

Because the LIEM is under-constrained, there are infinite possible solutions that satisfy the equality and inequality constraints. To choose a mean solution, and to determine uncertainty, within the possible solution space, we use a Markov Chain Monte Carlo (MCMC) sampling method (Kones et al., 2009; van den Meersche et al., 2009; van Oevelen et al., 2010), which has been shown to reconstruct unmeasured flows more accurately than the L_2_ minimum norm approach (Saint-béat et al., 2013; Stukel et al., 2012, 2018a). Implementation details are provided in the online supplement.

As a metric for discussing model results with respect to the approximation equations (i.e. the observations), we use the model-observation misfit relative to the model uncertainty: Σ = (*X*_*model*_−*X*_*obs*_)/*σ*_*obs*_. Here *X*_*model*_ is the model prediction, *X*_*obs*_ is the observed value, and *σ*_*obs*_is the standard deviation of the observed value. The square of this quantity (Σ^2^) is summed over all approximate equations yielding the solution cost function, and thus Σ is a proxy for disagreement between the LIEM and observations. Unless otherwise stated, LIEM solutions will be given as a range based on the mean solution for each cycle as well as the median value of all cycles. Displaying data in this way allows us to highlight inter-cycle variability. For value and uncertainty in all rate constraints, see Appendix A.

### 2.3. Analyses and model comparisons

#### 2.3.1. Indirect analysis

An indirect analysis permits investigation of the contributions of carbon between any two compartments through indirect linkages. By taking the normalized matrix of flows between compartments (G) and the identity matrix (I), the matrix (I-G)^−1^ provides all the indirect flow data (Kroes, 1977). In this way the contribution of the surface compartments to the deep ones can be ascertained even when no direct flows exist. For example, if the food chain was A → B → C; an indirect analysis would reveal that 100% of the flows to C go through A.

#### 2.3.2. Independent DVM estimates

A model to predict the export flux due to zooplankton DVM was recently published by Archibald et al. (2019), which adds a diel vertical migration module to the Siegel et al. (2014) ecosystem model. The Archibald et al. model parameterizes the export production based on NPP, size-fractionated grazing (i.e., protists and mesozooplankton), and the proportion of DVM mesozooplankton. The export production attributed to vertical migrators who defecate at depth is a function of total grazing, the gut clearance rate, and the proportion of zooplankton undergoing DVM (Eq. 2).

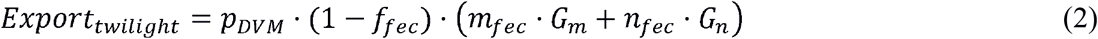

Where *p*_*DVM*_ is the fraction of mesozooplankton that undergo DVM, and *f*_*fee*_ is the fraction of fecal pellets produced by diel vertical migrators in the euphotic zone. *m*_*fee*_ and *n*_*fee*_ are the proportions of grazing that are exported by mesozooplankton and protistan zooplankton, respectively. *G*_*m*_ and *G*_*n*_ are the grazing rates for mesozooplankton and protistan zooplankton, respectively.

The respiration conducted by vertically migrating zooplankton can be calculated based on the metabolic efficiency, fraction of mesozooplankton undergoing DVM, and their grazing rate (Eq. 3).

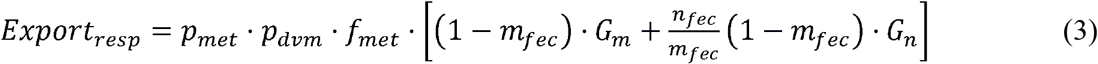

where *p*_*met*_ is the temperature dependent metabolic rate with ΔT the temperature difference between the mesopelagic and epipelagic and *p*_*met*_ = 2^(Δ*T*/10)^/(2^(Δ*T*/10)^ + 1). *f*_*met*_is the metabolic efficiency of the zooplankton, assumed to be 0.50. We calculated active transport from Equations 3 and 4 following Archibald, but using the CCE-optimized parameter set that Stukel et al. (2015) determined for the Siegel et al. (2014) model. The fraction of mesozooplankton undergoing DVM (*p*_*dvm*_) was calculated as described in Section 2.1.2. Fecal pellet production for meso- and microzooplankton were set to *m*_*fee*_ = 0.3 and *n*_*fee*_= 0.06 (Archibald et al., 2019), respectively.

Since the Archibald et al. model does not include mortality at depth as export and excludes any mesopelagic ingestion or excretion, the total export flux is the sum of Eq. 2 and 3. To compare with the LIEM presented here, a modified LIEM active transport flux will be calculated using the total active transport for mesozooplankton and subtracting mesopelagic mortality.

## 3. Results

### 3.1. *In situ* Ecosystem Observations

The locations for each study site were chosen to maximize the range of environmental conditions (Figure 2). Sea surface chlorophyll *a* (Chl *a*) varied from 0.2 – 1.5 mg Chl *a* m^−3^ with vertically-integrated primary productivity varying from 325 – 2314 mg C m^−2^ d^−1^. Productivity and biomass typically declined with distance from the Point Conception upwelling center. Most cycles were in water masses with steady or declining phytoplankton biomass (Figure 2D), with the exception of P0810-1. Sediment trap-derived carbon export at 100 m depth varied from 32 – 170 mg C m^−2^ d^−1^ (Figure 3C), with observed e-ratios (i.e. sediment trap export / ^14^CPP) ranging from 5% - 33%. Standing stock of zooplankton correlated positively with NPP and export (Spearman correlations of 0.36 and 0.40, respectively). Protistan zooplankton were responsible for grazing ~50% of NPP (Figure 3B) while mesozooplankton grazed, on average, ~30% of NPP with one exception (Figure 3E). The proportion of mesozooplankton biomass exhibiting DVM behavior ranged from 35% - 86% (median: 58%). Epipelagic bacterial production rates did not correlate with NPP but ranged from 22 – 400 mg C m^−2^ d^−1^ (Figure 3F), with the three lowest rates observed during the P0704 Cruise.

**Figure 2.**
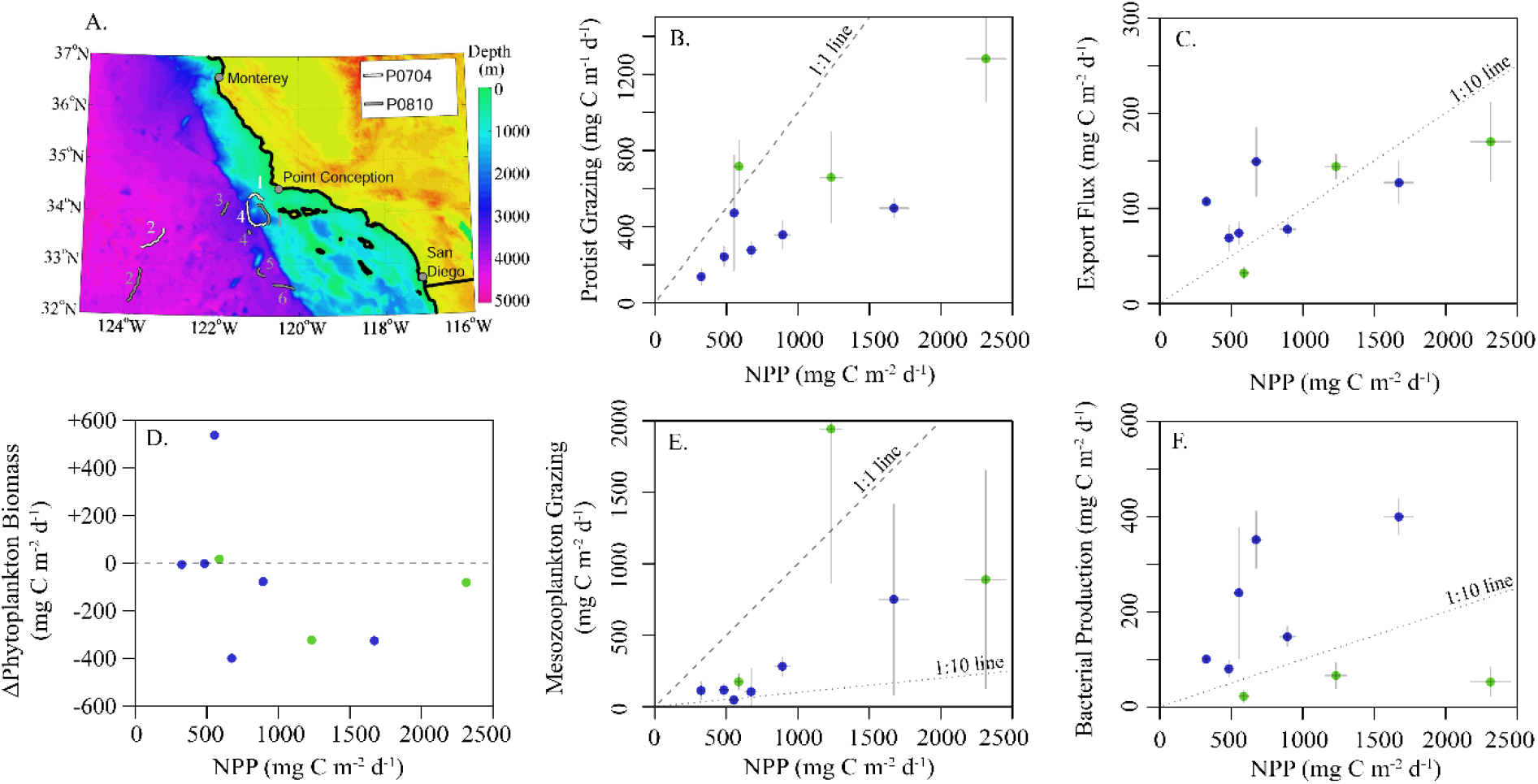
A) Bathymetric map of study region showing drift trajectories from each cycle. Numbers correspond to cycle for P0704 (white) and P0810 (grey). (B-F) Summary of *in situ* observations plotted against NPP: (B) protistan grazing, (C) export flux from sediment trap at 100 m, (D) observed rate of change of phytoplankton biomass, (E) mean mesozooplankton grazing and (F) epipelagic bacterial production. Values are colored by cruise (P0704 = green, P0810 = blue). Dashed lines for reference slopes of 1:1, 1:10, or no change as indicated and error bars are ±1 SD.

**Figure 3.**
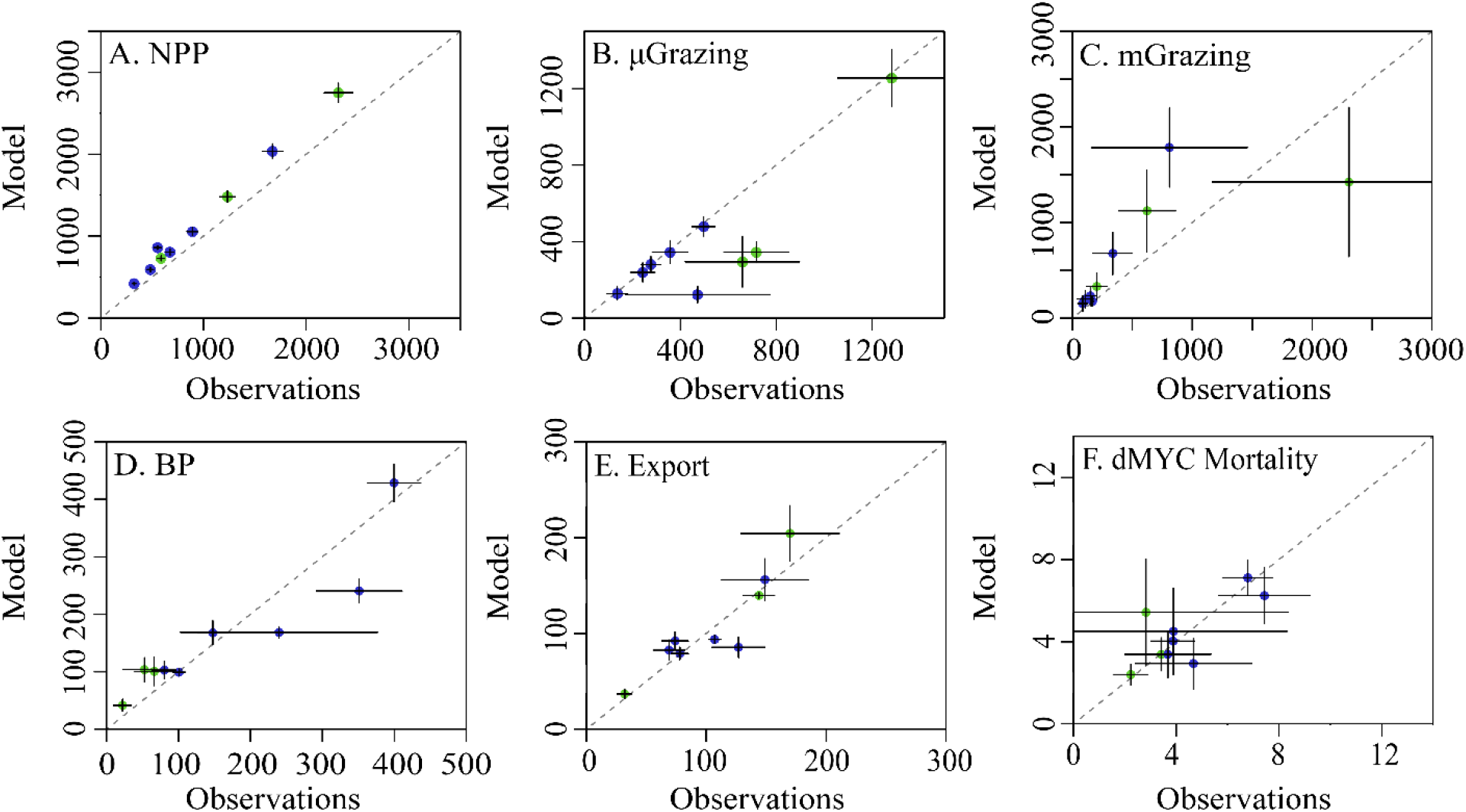
Model-observation comparisons for selected measurements: (A) net primary productivity, (B) protistan zooplankton grazing, (C) mesozooplankton grazing, (D) epipelagic bacterial production, (E) sediment trap carbon export (@ 100m), and (F) non-vertically migrating mesopelagic nekton mortality. Cruises are denoted by color (P0704 = green, P0810 = blue). Dashed line is 1:1 and error bars show 1 SD of uncertainty.

### 3.2. Model-Observation Mismatch

The LIEM solutions consistently show general agreement with all *in situ* observations except for modeled NPP, which is elevated by 18% - 56% (median: 22%) from ^14^CPP estimates (Figure 3A), or 3.0 – 9.3 Σ (median: 3.6 Σ). This degree of misfit corresponds to 18% – 82% (median: 46%) of the total model-observation misfit. Model agreement with the sediment trap was high (−33% – 25%; Figure 3E) with a modeled e-ratio (i.e. sediment trap export / NPP) of 5% – 35% (median: 14%), which compares well to the observed e-ratio of 5% – 33% (median: 11%). Modeled protistan grazing rates and mesozooplankton grazing rates were reasonably close to observations (Figure 3B). Modeled microzooplankton (MIC) grazing was lower than observed for cycles P0704-2 (−2.8 Σ) but agreed reasonably well (−1.5 – +0.1 Σ) for the other cycles (Supplemental Figure 1). For P0704-1, mesozooplankton grazing rates were lower than observations for SMZ (−1.8 Σ), total non-DVM grazing (−1.8 Σ) and for vmSMZ grazing (−1.6 Σ). This cycle exhibited declining phytoplankton biomass (−322 mg C m^−2^ d^−1^) and very high zooplankton grazing rates compared to the other cycles. This water parcel may have been in a declining bloom stage where observed grazing rates were unsustainable. Model-data agreement among the seven nekton-related observations (e.g. Figure 3F) was satisfactory (|Σ| < 1) except for P0810-1, which showed reduced vertically migrating nekton activity relative to estimates (vmMYC epipelagic respiration: −1.5 Σ, vmMYC mesopelagic respiration: −1.7 Σ, and vmMYC mesopelagic mortality: −1.1 Σ). This cycle was along the edge of an anti-cyclonic eddy, where lateral gradients were likely high.

### 3.3. Epipelagic Ecosystem Model

According to the LIEM, phytoplankton respired 18% - 39% (median: 30%) of GPP, lost 14% - 26% (median: 18%) as DOC, lost 2% - 42% (median: 6%) to non-grazer mortality and the remaining 5% - 54% (median: 45%) was grazed by zooplankton. Modeled NPP ranged from 421 mg C m^−2^ d^−1^ to 2750 mg C m^−2^ d^−1^ (median: 861 mg C m^−2^ d^−1^). The LIEM suggested that protists and mesozooplankton had relatively similar grazing impacts on phytoplankton across all cycles, although the proportional role was greater for mesozooplankton in coastal regions and greater for protists during oligotrophic conditions. Between 14% and 47% (median: 33%) of NPP was grazed by protistan zooplankton (MIC + HNF) and 18% - 96% (median: 45%) by mesozooplankton (SMZ + vmSMZ + LMZ + vmLMZ). We note that protistan grazing rates normalized to NPP are slightly depressed relative to observations since model NPP was higher than observations while protistan grazing generally matched the observations (Figure 3; Supplemental Figure 1). 57% - 82% (median: 74%) of mesozooplankton grazing was by small mesozooplankton (SMZ + vmSMZ). Vertically migrating mesozooplankton were responsible for 52% - 89% (median: 63%) of total mesozooplankton grazing, 58% - 85% (median: 77%) of which was done by small mesozooplankton (i.e. vmSMZ grazing / total vm grazing).

Mortality relative to ingestion for mesozooplankton was similar for the different epipelagic mesozooplankton (i.e. SMZ, LMZ, vmSMZ, and vmLMZ): SMZ: 24% - 25%, vmSMZ: 23% - 25%, LMZ: 22% - 25%, and vmLMZ: 24% - 27%, as was fecal pellet production (between 30% and 40% of ingestion).

Overall, 19% - 44% (median: 29%) of NPP was transferred from the epipelagic to the mesopelagic with 3% - 8% (median: 5%) of NPP leaving the epipelagic through higher trophic levels (SAR + vmMYC). Gravitational settling and subduction of POC accounted for 14% - 78% (median: 55%) of epipelagic export, while 18% - 84% (median: 41%) was through active transport of DVM mesozooplankton (vmSMZ + vmLMZ). Vertically migrating myctophids (vmMYC) transferred 2% - 6% (median: 4%) of total export. Section 3.4 provides a more detailed description of export production.

The gross growth efficiencies (GGE) for each type of organism are shown in figure 4A. Overall, BAC GGE was 7% - 29% (median: 25%) with an upper bound set to 30%. Notably, BAC GGE differed based on cruise, with P0704 cycles ranging between 8% - 13% and P0810 ranging between 23% - 29%. MIC GGE was 35% - 38% (median: 37%), and HNF GGE ranged from 32% - 35% (median: 33%), which is slightly higher than typical estimates of protistan zooplankton GGE (Straile, 1997) although reported variability is high (Steinberg and Landry, 2017). Epipelagic mesozooplankton GGE were consistently above 20%.

**Figure 4.**
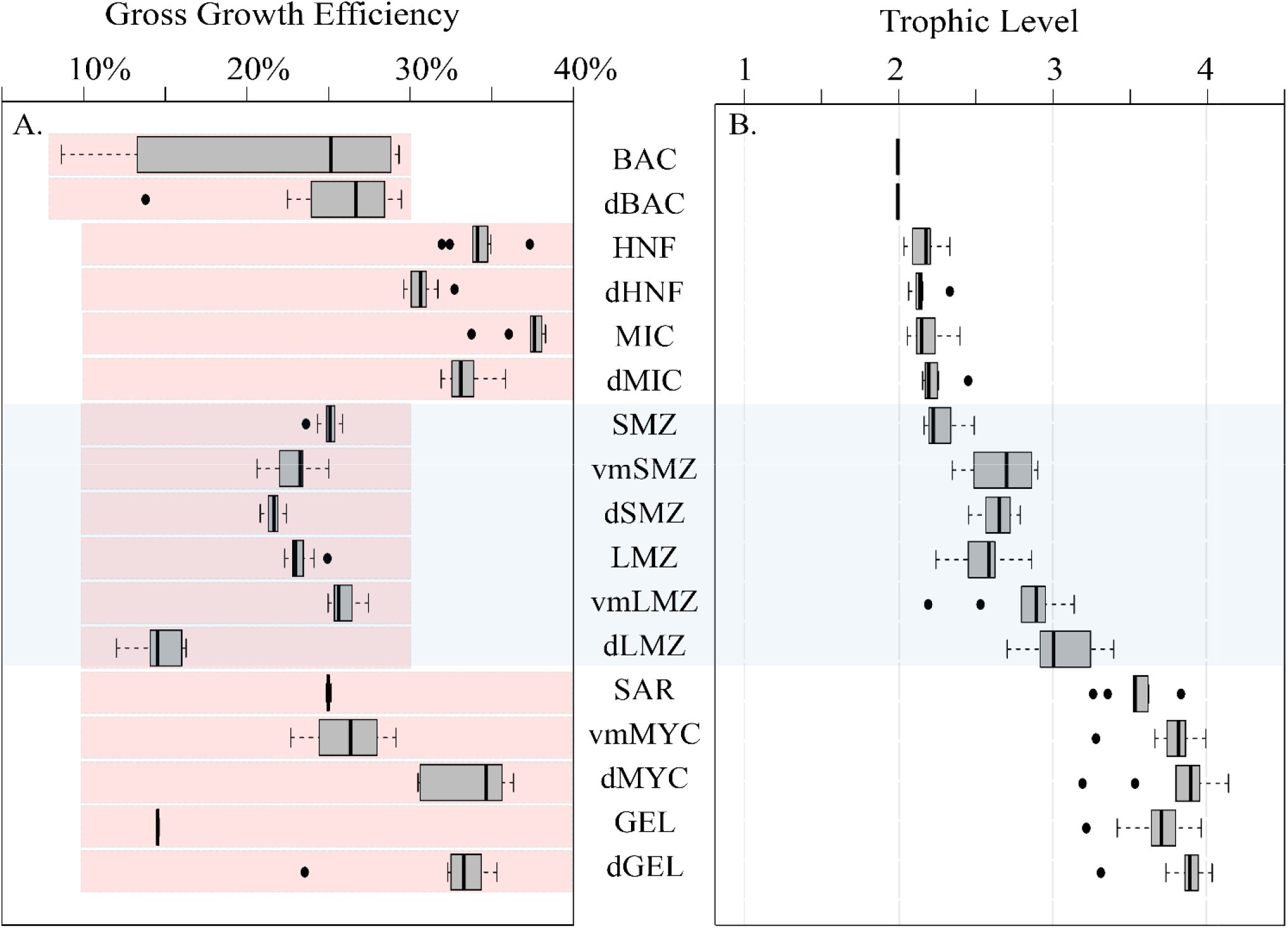
(A) Box and whisker plot of GGE for organisms in the LIEM. Red shaded boxes indicate the permitted range of values constraining the LIEM. (B) Box and whisker plot of trophic levels for each zooplankton assuming detritus and primary productivity are trophic level 1, and bacteria are trophic level 2. Box and whisker plots show inter-quartile range and 95% C.I. as determined using the mean solutions for each cycle. For reference the mesozooplankton compartments are shaded across both figures. Abbreviations are explained in Table 2.

#### 3.3.1 Trophic Level & Diets

Trophic levels for each organism (Figure 4B) were calculated by assuming that primary productivity, detritus and DOC were at trophic level 1. Trophic level indices were not affected by the overall cycle productivity (i.e. NPP), time of year, or by nutrient regime. The trophic level of small epipelagic mesozooplankton (SMZ) ranged from 2.2 to 2.5 (median: 2.2) and large mesozooplankton (LMZ) ranged from 2.2 to 2.9 (median: 2.6). The SAR trophic level was 3.3 – 3.8 (median: 3.5), and vmMYC was similar at 3.3 – 4.0 (median: 3.8). Modeling these higher trophic levels is important for structuring the ecosystem, and the nekton trophic levels found here are consistent with findings from ^15^N amino acid studies (Choy et al., 2015).

The modeled mesozooplankton ingestion can be classified into four distinct dietary types: (1) Herbivory = phytoplankton diet, (2) Protistivory = protistan zooplankton diet, (3) Detritivory = detrital diet (i.e. SDT or LDT), and (4) Carnivory = mesozooplankton diet. Using this partitioning, the relative contributions of each dietary component were assessed for large and small vertically migrating mesozooplankton compartments (Figure 5). The largest proportion of the diet for resident epipelagic mesozooplankton (i.e. SMZ & LMZ) was balanced between herbivory (19% - 57% median: 40%) and protistivory (26% - 59% median: 40%). Detritivory was 9% - 21% (median: 13%) of total diet. Inter-cycle variability in carnivory was low for resident epipelagic mesozooplankton and contributed 6% - 8% (median: 6%) of their diet.

**Figure 5.**
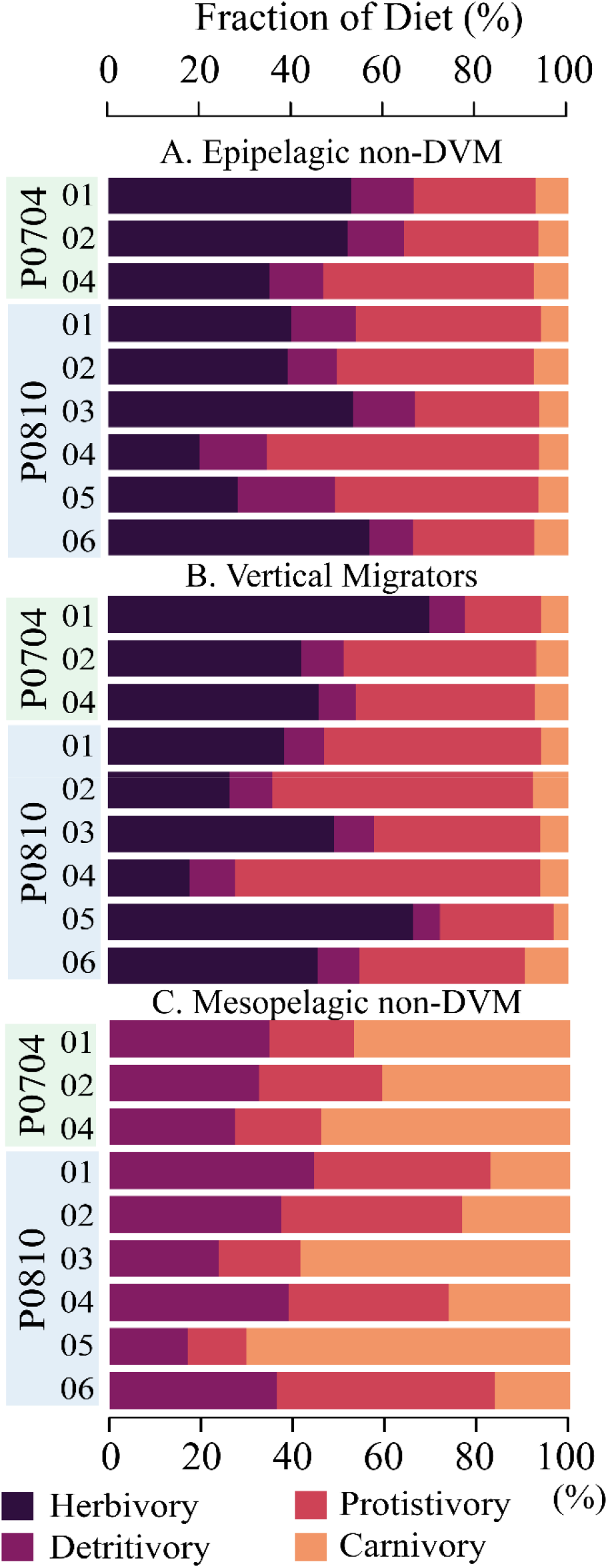
Composition of diet for (A) epipelagic, non-DVM mesozooplankton, (B) vertically migrating mesozooplankton, and (C) mesopelagic, non-DVM mesozooplankton. Diet is partitioned into herbivory (darkest), protistivory, detritivory, and carnivory (lightest). Cycles are as indicated.

Comparing the LIEM solutions between the nutrient limited and upwelling cycles, we found that large mesozooplankton grazing increased from 9% - 16% (median: 13%) in the nutrient limited cycles to 22% - 65% under upwelling conditions (median: 30%) of NPP, although the overall diets of the mesozooplankton did not systematically change with nutrient condition.

### 3.4. New Production, Export and DVM

Total export ranged from 163 - 707 mg C m^−2^ d^−1^ (median: 282 mg C m^−2^ d^−1^) with distinctly elevated values associated with upwelling cycles (Figure 6A). The fraction of export attributed to mesozooplankton DVM (vmSMZ + vmLMZ) covaried with nutrient regime: mesozooplankton active transport contributed 14% - 37% of total export under nutrient limited conditions and 44% - 84% under upwelling conditions (Figure 6B). There was not a significant relationship (p < 0.1) between the total export efficiency (i.e. total export / NPP) and NPP (Figure 6C).

**Figure 6.**
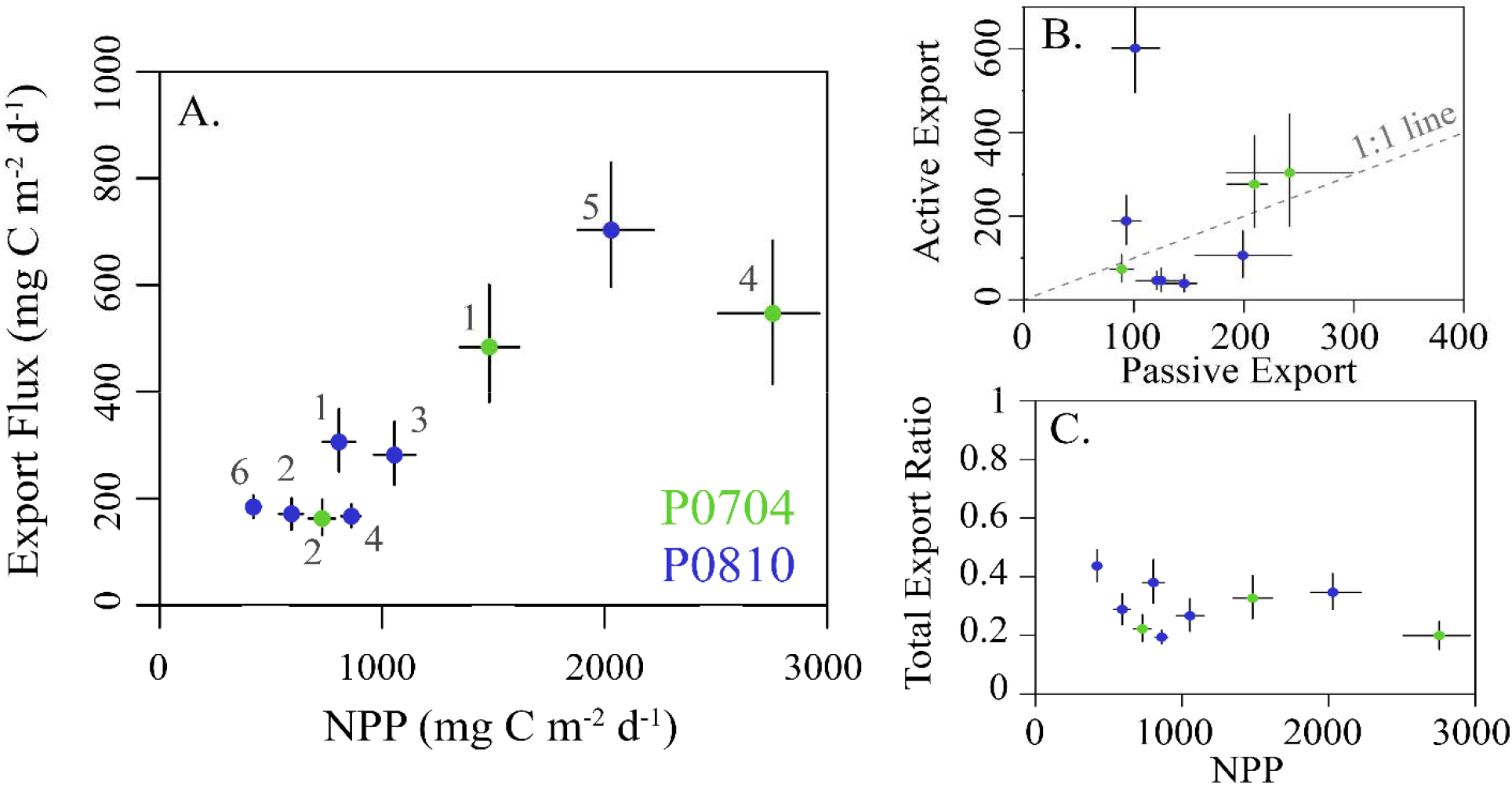
(A) The sum of both passive and active carbon export flux from the epipelagic plotted against NPP. Cruises are color coded and error bars show the 95% CI for each value. (B) The total active flux due to DVM verses passive flux for each cycle (as in A). Dashed 1:1 line for reference. (C) The total export ratio (i.e. total epipelagic export / NPP) plotted against NPP and colored as in (A). All values are in mg C m^−2^ d^−1^.

For vmSMZ, 77% - 80% (median: 80%) of their respiration took place in the epipelagic, along with 67% - 87% (median: 85%) of their DOC excretion. This is consistent with the suggestion that mesozooplankton respiration and excretion are elevated in the warmer epipelagic waters (Ikeda, 1985) and where activity is highest.

The fate of active export flux is important for understanding the ecological impact of this carbon supply. Within the mesopelagic, mesozooplankton respired 11 – 104 mg C m^−2^ d^−1^ (median: 33 mg C m^−2^ d^−1^) and excreted 7 – 116 mg C m^−2^ d^−1^ (median: 20 mg C m^−2^ d^−1^; Figure 7A). Predation on vertically migrating mesozooplankton accounted for a loss of 23 – 352 mg C m^−2^ d^−1^ (median: 59 mg C m^−2^ d^−1^) in the mesopelagic. Mesozooplankton fecal pellet production in the mesopelagic was 8 – 29 mg C m^−2^ d^−1^ (median: 13 mg C m^−2^ d^−1^). Resident mesopelagic mesozooplankton were the dominant mortality term for the vertically migrating mesozooplankton (Figure 7B).

**Figure 7.**
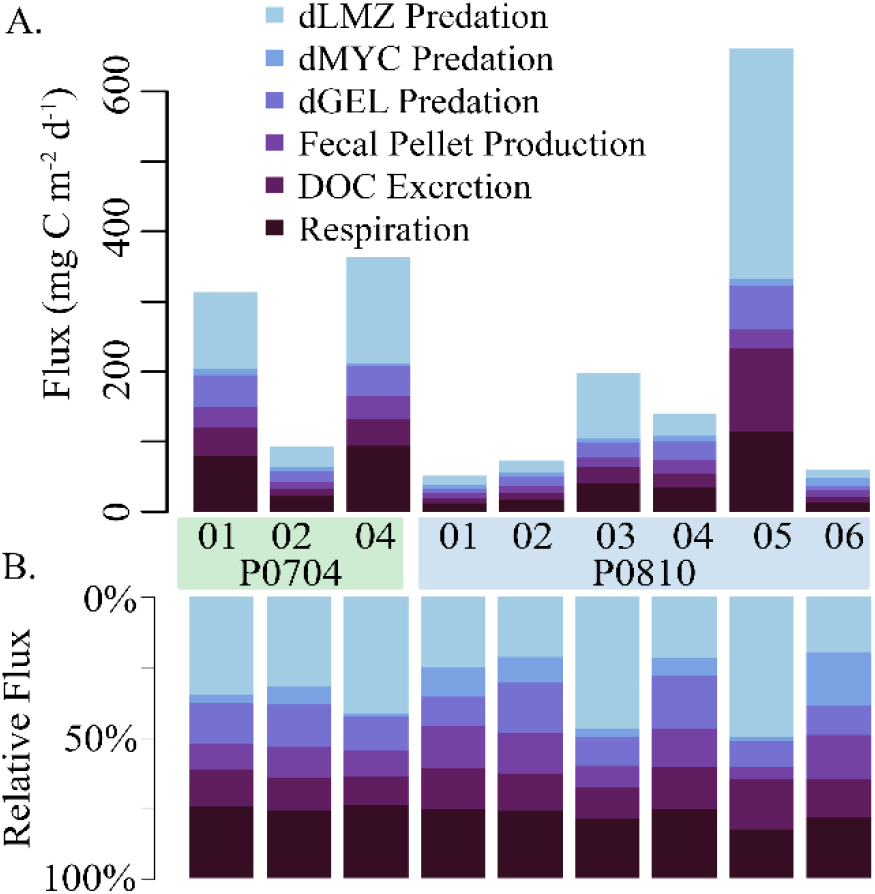
The (A) net and (B) relative fate of vertically migrating mesozooplankton within the mesopelagic. Loss terms are color coded, and cruise and cycle are as shown. Abbreviations are explained in Table 2.

### 3.5. Mesopelagic Ecosystem

Deep bacteria (dBAC) made up 6% - 30% (median: 11%) of the mesopelagic protistan zooplankton diet with the remainder supplied by detritus/fecal pellets. Mesopelagic mesozooplankton (i.e. dSMZ & dLMZ) had a more variable diet than the epipelagic mesozooplankton (Figure 5), with detritivory ranging from 17% - 43% (median: 39%) of their diet, protistivory at 14% - 51% (median: 30%) and carnivory at 10% - 68% (median: 33%).

Systematic increases in trophic level between the epipelagic and mesopelagic resident zooplankton and nekton were observed (Figure 4). The trophic level of epipelagic microzooplankton (MIC) was 2.0 – 2.3 while dMIC was 2.3 – 2.5. Similar increases between the epipelagic and mesopelagic were observed for mesozooplankton, where SMZ had a trophic level of 2.2 – 2.5 (median: 2.2) dSMZ had a trophic level of 2.5 - 2.8 (median: 2.6). Likewise, dLMZ trophic levels were elevated by ~0.4 relative to LMZ. The trophic level of dMYC (3.2 – 4.1) was more variable than the other micronekton (e.g. vmMYC: 3.5 – 4.0), illustrating a greater variability in diet.

Mesopelagic respiration is a useful diagnostic loss term for determining which organisms are responsible for the mesopelagic carbon demand (Supplemental Figure 2). Mesopelagic bacteria accounted for the largest proportion of mesopelagic respiration (31%-41% median: 34%). High respiratory term for mesopelagic bacteria was found despite relatively high GGE for these organisms (median 26%, Figure 4A). Mesopelagic protistan zooplankton and resident mesozooplankton were responsible for 14% - 30% (median: 25%) and 14% - 24% (median: 15%), respectively. Resident gelatinous predators and myctophids are responsible for 4% - 8% o mesopelagic respiration combined. The proportion of export due to active transport covaried with resident mesopelagic respiration (Figure 8A), illustrating the coupling between active transport and mesopelagic activity in the LIEM. The effect of higher active transport relative to total export can be shown with an indirect analysis where the relative contribution of carbon from epipelagic detritus (i.e., a passive transport proxy) and vertically migrating mesozooplankton (i.e., an active transport proxy) in the diet of each organism can be measured. Indirect flux analyses show that a higher proportion of the carbon consumed by mesopelagic bacteria, protists, and mesozooplankton originated from passive rather than active transport (Figure 8B). However, mesopelagic nekton (dMYC) were predominantly supported by carbon derived from active transport pathways.

**Figure 8.**
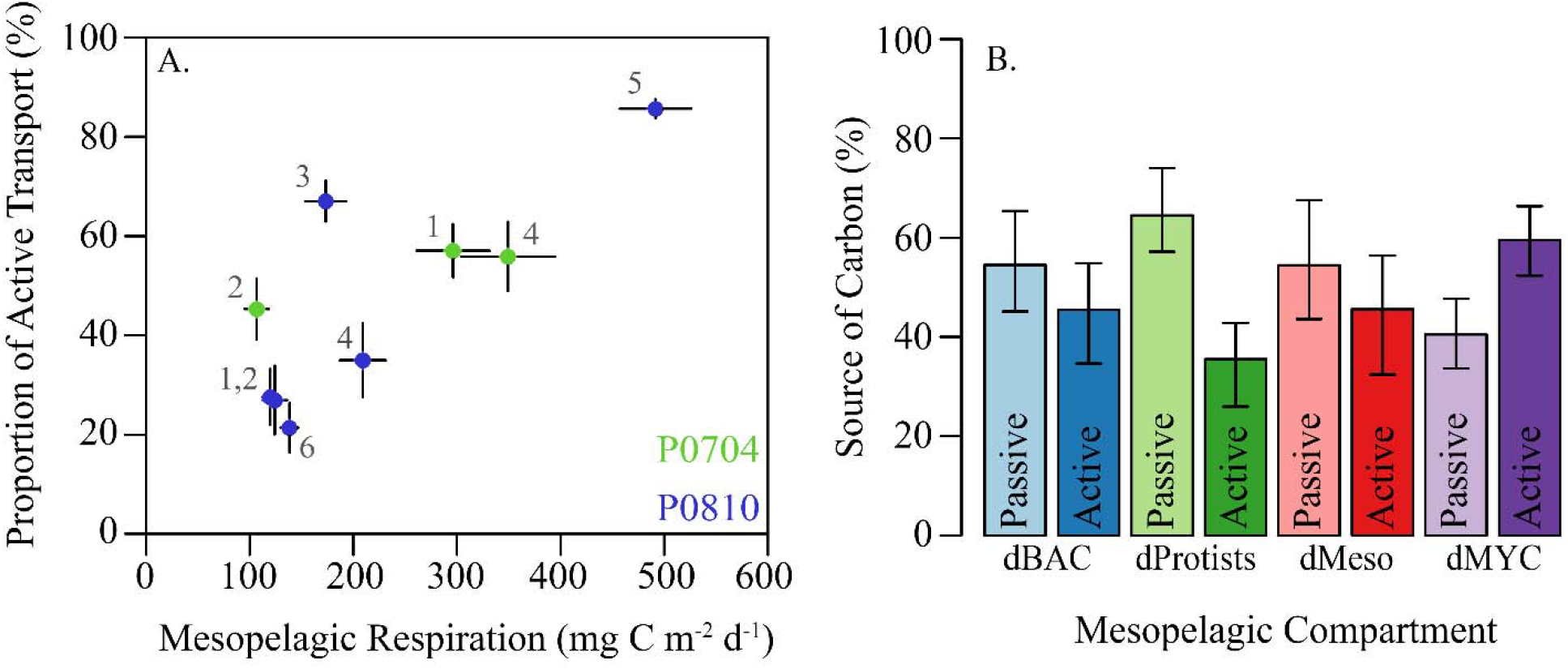
(A) Proportion of active transport relative to total export verses total mesopelagic respiration by residents (i.e. dBAC, dHNF, dMIC, dSMZ, dLMZ, dMYC, dGEL). Cruises are colored and cycles are as shown. (B) Relative proportions of carbon demand supplied by passive or active flux for the indicated mesopelagic groups. Source was calculated using indirect flux analysis (Section 2.3.1) to determine the indirect contribution epipelagic detritus (passive) and vertically migrating mesozooplankton (active). Error bars are ±1 SD.

## 4. Discussion

### 4.1. Diel Vertical Migration & Active Transport in the CCE

In contrast to common assumption about the processes driving the biological pump, our results suggest that active transport may be as, if not more, important than sinking particle flux. We found that active transport (mesozooplankton and fish combined) was responsible for 39 - 606 mg C m^−2^ d^−1^ (median: 107 mg C m^−2^ d^−1^), corresponding to 21% - 86% of total export to the mesopelagic, while sinking particles contributed 14% - 79%. This finding is not directly forced by an *a priori* assumption of the importance of active transport. Indeed, we placed no direct constraint on the amount of mesozooplankton mortality in the mesopelagic, and the minimum constraints on basal metabolism by zooplankton in the mesopelagic (Stukel et al., 2013) implied that active transport could have been as low as 2% – 40% of sinking flux (median: 18%). Nevertheless, the importance of active transport was a robust result of the inverse analyses. For P0810-6, the cycle with the lowest relative contribution of active transport to total export (21%), the total flux was 184 ± 23 mg C m^−2^ d^−1^ (95% CI) and active transport was 39 ± 21 mg C m^−2^ d^−1^ (95% CI). This cycle was oligotrophic and had the lowest ^14^CPP measurements of any cycle on the two cruises. In contrast, cycle P0810-5 had the highest relative contribution of active transport (86% ± 4% of total export at the 95% CI). P0810-5 was on the coastal (i.e. high biomass) side of a strong frontal feature with high rates of primary productivity and large standing stocks of zooplankton.

Although these rates of active transport are higher than reported in many studies, they are fully consistent with mesozooplankton community dynamics in the CCE. The model suggests that total epipelagic mesozooplankton consumption on phytoplankton, protists, detritus, and other mesozooplankton ranged from 361 - 2966 mg C m^−2^ d^−1^ (median: 1006 mg C m^−2^ d^−1^). Vertically stratified day-night net tows showed that 35% - 86% (median: 57%) of the mesozooplankton community was vertically migrating to depth each day and that most of these vertical migrants were copepods and euphausiids (Stukel et al., 2013). Our model results indicate that only 20% - 23% of respiration and 16% - 34% of excretion by vertical migrants occurred at depth. None of these assumptions are particularly aggressive. Furthermore, our results (Figure 9) are consistent with estimates of DVM in the zooplankton derived from the model of Archibald et al. (2019), if specific dynamics of the CCE are taken into account (e.g., zooplankton consume nearly all of NPP, Landry et al. 2009; microphytoplankton are negligible contributors to sinking flux, Stukel et al. 2013). Our estimates of the total export ratio 19% - 44% are also consistent with typical *f*-ratio estimates (new production to total export) in our study region, which varied from 0.23 to 0.40 (Krause et al., 2015). Our results thus do not arise from unusual parameterizations but instead may reflect the fact that estimates of active export using standard metabolism calculated from Ikeda et al. (1985; 2001) may be conservative underestimates.

**Figure 9.**
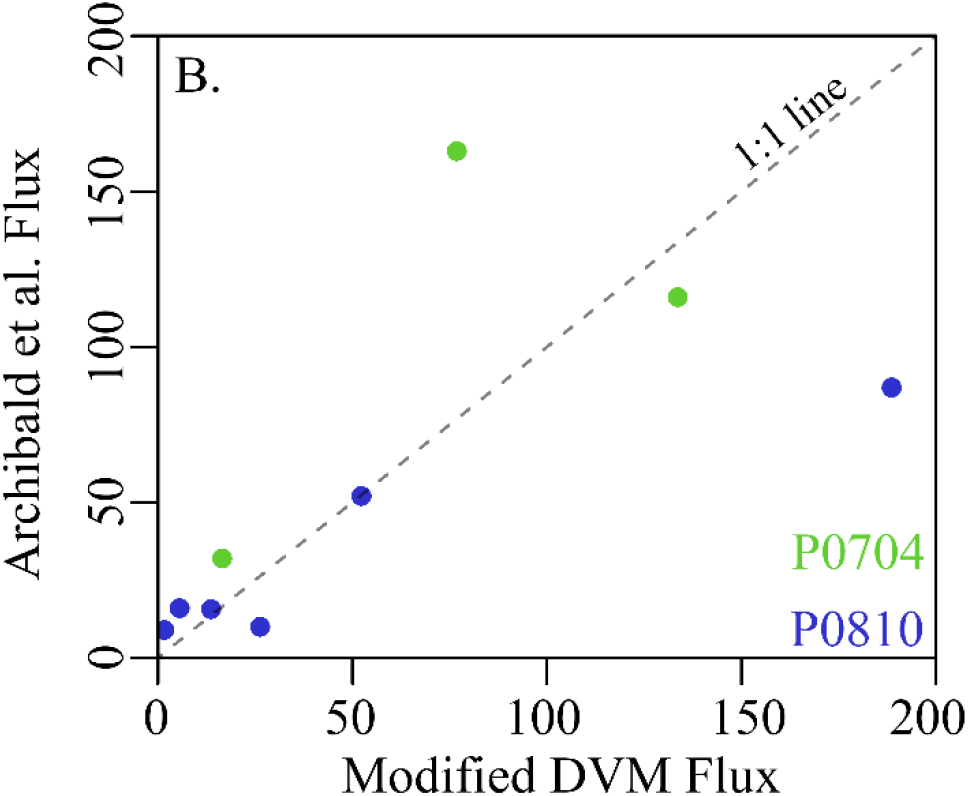
Comparison between modeled mesozooplankton DVM flux without mesopelagic mortality and the predicted flux from Archibald et al. (2019) with CCE-specific parameterization following Stukel et al. (2015). Cruises are as colored (P0704 = green, P0810 = blue) and dashed line is a 1:1 reference line. Fluxes are shown in mg C m^−2^ d^−1^.

Our results also reflect realistic coupling between the epipelagic and mesopelagic communities. Model results suggested that the carbon demand was equal to <1% - 4% (median: 1.1%) of NPP for mesopelagic fish, 1% - 7% (median: 3%) of NPP for predatory gelatinous zooplankton, 8% - 22% (median: 14%) of NPP for resident mesopelagic zooplankton, and 6% - 19% (median: 11%) of NPP for mesopelagic bacteria. These mesopelagic carbon demands must be met by carbon flux from the surface layer, the most likely sources of which are sinking particle flux (which we experimentally measured using two independent approaches) and active transport. While it is possible that both sediment traps and ^238^U-^234^Th disequilibrium underestimated sinking carbon flux, the inverse analysis offers compelling evidence that active transport is more likely to support mesopelagic fish and gelatinous predator communities. Although sinking particles can efficiently support bacterial production (as they are likely directly colonized by particle-attached bacteria), many fish and gelatinous zooplankton are predators that more likely feed on living organisms than on the sinking fecal pellets that typically dominate particle flux in the CCE. For these planktivorous organisms, sustaining their metabolism through a food chain supported by sinking particles would likely require one (if not more) trophic levels to separate them from the export source, depending on whether the sinking particles are consumed by filter- or flux-feeding zooplankton or by microbes (Stukel et al., this issue). Thus, sustaining the high carbon demand of mesopelagic myctophids with sinking particles requires substantially more total carbon flux than does sustaining it via active transport of the myctophids’ prey.

Mesopelagic sources of mortality have implications for the fitness of vertical migrators. It is often assumed that DVM is ecologically advantageous when the costs associated with not feeding during the day and actively swimming to depth are offset by the benefits of reduced predation pressure and/or reduced metabolism at colder mesopelagic temperatures (Bianchi et al., 2013; Hansen and Visser, 2016; Morozov and Kuzenkov, 2016). Our model suggests that mortality normalized to ingestion is similar across all mesozooplankton compartments and across a wide range of ecosystem states (SMZ: 24% - 26%, LMZ: 22% - 25%, vmSMZ: 21% - 25%, vmLMZ: 25% - 27%, dSMZ: 21% - 23%, dLMZ: 19% - 23%). Even though vmSMZ experience similar predation to SMZ and dSMZ, approximately half of the predation on vertically migrating zooplankton takes place in the mesopelagic, thereby transferring carbon to depth despite the fact that their excretion and respiration occur primarily in the epipelagic.

The comparable mortality experienced by vertically-migrating mesozooplankton in the mesozooplankton may seem counterintuitive in light of extensive research suggesting that the adaptive advantage of DVM may be to reduce predation (Bandara et al., 2018; Ohman and Romagnan, 2016). However in the CCE, it is not particularly surprising when the large abundances of myctophids, gonostomatids, and other mesopelagic fish are considered. Davison et al. (2013, 2015) demonstrated high biomass of these fish comprising both vertically-migrating and mesopelagic resident communities. These organisms may thus face as high, if not higher, predator abundance at mesopelagic depths than in the epipelagic, although colder temperatures and reduced irradiance may diminish predation rates at depth. DVM may remain advantageous as a lifestyle because if these organisms were present at the surface during the day then they might experience substantially greater predation than in the mesopelagic.

### 4.2. Sensitivity Analysis and Ecological Connections

The ecosystems generated in the 9 model runs were as varied as the cruise measurements: including observations from dynamic coastal blooms to quiescent oligotrophic communities. All 9 cycles had significantly elevated NPP compared to the observed ^14^CPP (Figure 3; Supplemental Figure 1) with 95% CI from the MCMC random walk. Whether this result can be considered a model bias or is derived from possible systematic differences between ^14^CPP and true net primary production (Marra, 2009; Milligan et al., 2014; Minas et al., 2002; Peterson, 1980) is not known. To test the model’s sensitivity to the misfit with ^14^CPP, the LIEM was rerun assuming that ^14^CPP uncertainty was 1/10^th^ of the actual estimated uncertainty. The model-observation misfit increased by nearly 2.5x with small mesozooplankton grazing rates, myctophid metabolic estimates, and sediment trap export all reduced by ~2 Σ relative to the standard model run. This result shows that the model needed to increase NPP in the standard model run in order to match the observed mesozooplankton grazing rate and myctophid metabolic requirements.

Because bacterial activity in the mesopelagic was not measured we made conservative assumptions about the possible upper and lower bounds for bacterial production. For the minimum bound on mesopelagic BP, we chose a conservative power-fit slope of −1.47 (Yokokawa et al., 2013). This resulted in model-determined mesopelagic bacterial carbon demand that may have been lower than true *in situ* values. Other reported values for the attenuation of BP in the mesopelagic include slopes of *α* = 1.15 (Tanaka and Rassoulzadegan, 2004) and 1.03 (Gasol et al., 2009), which would result in 25% and 36% higher estimates of mesopelagic BP, respectively. When the minimum mesopelagic bacterial production estimates were doubled (a = 0.64; Eq. 1), the model responded by increasing NPP by +2% (inter-cycle median) and total export flux by 11%. Since passive particle flux is constrained by observations, passive flux increased by 0% - 12% (median: 4%) while active transport by mesozooplankton increased by 0% - 56% (median: 26%). Active transport by nekton was also elevated (0% - 14%, median: 10%). Model-observation misfit increased by an average of 17% with notable changes in NPP (+0.42 Σ), sediment trap flux (+0.34 Σ) and Thorium-234 flux (+0.22 Σ).

The results were also robust to changes in other observations. When the nekton metabolic estimates were halved, export by vmMYC was reduced by 51% (inter-cycle median), a change of < 5 mg C m^−2^ d^−1,^ while other forms of export were unchanged. Increasing the upper limit of mesozooplankton GGE from 30% to 40% led to a ~20% increase in mesozooplankton active transport and no change in nekton-derived active flux or passive flux.

### 4.3. Linear Inverse Models

LIEMs are a powerful tool for assimilating diverse *in situ* measurements and constraints with a food web perspective. The use of a two-layer model (Jackson and Eldridge, 1992) is particularly powerful because it allows information from the mesopelagic to constrain epipelagic food web flows and vice versa. Compared to most previously published LIEMs, the model presented here includes many more *in situ* rate measurements, made possible by the suite of contemporaneous rate measurements made during quasi-Lagrangian experiments. For comparison, many LIEMs are constrained by fewer rate measurements (Dubois et al., 2012; Sailley et al., 2013; van Oevelen et al., 2012) and therefore rely much more heavily on greater than/less than constraints derived from biomass measurements, leading to correspondingly higher uncertainty. This highlights a need for studies that simultaneously quantify the activity of many different plankton functional groups.

Since a LIEM is fundamentally a data-regression technique, our results are emergent from (A) our observations, (B) the conservative assumptions used (e.g. GGE), and (C) the ecosystem structure. Thus, we believe the resulting model solutions to be descriptive of the dominant *in situ* processes in the CCE LTER study region. However, it is important to note that there was large uncertainty associated with some model flows, and that this uncertainty could be quantified using the MCMC approach (Supp. Table 2). We thus highly recommend the MCMC approach (Kones et al., 2009; van den Meersche et al., 2009), which has been shown to more robust in its ability to recover ecosystem rates (Saint-béat et al., 2013; Stukel et al., 2012). Even more important is its ability to generate confidence intervals that realistically represent the uncertainties in model outputs with respect to both measurements and under-determinacy of the model. For instance, on cycle P0810-6, we found that the 95% confidence interval for HNF ingestion of detritus was 5 - 127 mg C m^−2^ d^−1^, providing no real knowledge of whether or not this connection was an important part of the ecosystem. However, for Cycle P0810-5, we found that mesopelagic mesozooplankton predation on small vertical migrators was 233 - 423 mg C m^−2^ d^−1^ (95% CI) and hence have a high degree of confidence that this flow was substantial at this location. Investigation of the confidence intervals can thus inform which conclusions can be considered robust. Developing even better-resolved ecosystem models likely requires incorporation of more diverse measurement types, such as ^15^N isotopic data (Stukel et al., 2018a).

### 4.4. The biological carbon pump and mesopelagic flux attenuation

Reports of active transport by vertically migrating biota have long suggested that these organisms can transport a globally significant amount of carbon to depth. However, most early studies suggested that active transport was substantially less important than passive flux of sinking particles (Davison et al., 2013; Morales, 1999; Steinberg and Landry, 2017). At the oligotrophic BATS station off Bermuda, Dam et al. (1995) found that respiration by mesozooplankton augmented the passive carbon flux at 150 m by 18% – 70%. Also at BATS, Steinberg et al. (2000) reported a significant vertical transfer of nitrogen by zooplankton, including dissolved organic nitrogen (DON). In fact, vertical migrators were found to perform 15% - 66% of the total nitrogen transport. Hansen and Visser (2016) estimated that across the North Atlantic active transport by mesozooplankton may constitute 27% of total export. In addition to zooplankton, vertical migrations by micronekton can also lead to significant export fluxes (Angel and Pugh, 2010; Davison et al., 2013; Hernandez-Leon et al.). Using biomass estimates and metabolic relationships, Davison et al. (2013) found micronekton contributions of 22 – 24 mg C m^−2^ d^−1^ (or 15% - 17% of estimated passive export) in the northeast Pacific. In the North Pacific Subtropical Gyre, Al-Mutairi and Landry (2001) estimated that active transport due to zooplankton respiration was responsible for carbon flux equal to 18% of passive flux. Using a conservative approach, Longhurst (1990) estimated that active export by zooplankton DVM was 13% - 58% that of passive flux when accounting for respiration alone in subtropical waters, which is similar to our results where the LIEM suggests that mesozooplankton respiration at depth is 9% – 113% (median: 34%) that of passive export. Global modeling estimates have suggested that active transport may be responsible for 14% (Archibald et al., 2019) or 15 to 40% (Bianchi et al., 2013) increases in carbon export relative to sinking particles alone. More recent results, have been suggesting increased importance for active transport, potentially rivaling that of passive flux. In the Costa Rica Dome, a region with high mesozooplankton biomass like the CCE, Stukel et al. (2018b) identified active transport by zooplankton DVM as responsible for 21-45% of total euphotic zone export. Hernández-León et al. (this issue) found that active transport was equal to one quarter of passive flux in oligotrophic regions, but was 2-fold higher than passive flux in eutrophic areas of the tropical and subtropical Atlantic. Our results that total active transport (zooplankton and nekton) may be responsible for 18% - 84% (median: 42%) of total carbon export in the CCE are thus somewhat higher than found in most studies, but not consistent with recently published values for high zooplankton biomass regions. Furthermore, our results are in line with other biogeochemical and ecological expectations (e.g., mesopelagic carbon demand, euphotic zone new production, mesozooplankton energy partitions). We thus suggest that active transport in high biomass regions may be more important, in fact, than some previous studies suggest, and we recommend focused research to investigate the potentially conservative assumptions made in previous studies that rely on standard (rather than active) estimates of zooplankton metabolic rates.

Within the mesopelagic, zooplankton also play an important biogeochemical role in the attenuation of particle flux (Buesseler and Boyd, 2009; Steinberg et al., 2008; Stukel et al., this issue) and in effecting elemental cycling (Kiko et al., this issue; Robinson et al., 2010). Our results suggest that mesozooplankton detritivory accounted for the consumption of 57% - 71% of sinking particles from the epipelagic, with bacterially-mediated remineralization of the majority of the remainder (i.e. mesopelagic export efficiency is < 10%). Notably, 3 of the 4 cycles with the lowest proportion of detritivory and the largest proportion of carnivory in the resident mesopelagic zooplankton were during upwelling cycles. This is the opposite of the findings of Wilson et al. (2010), who observed increases in fatty-acid biomarkers associated with carnivory at station Aloha relative to K2 and attributed the increase to the lower primary productivity at station Aloha. Our result that zooplankton rely more heavily on carnivory in the mesopelagic agrees with fecal pellet characteristic analyses and fatty acid biomarkers measured by Wilson et al. (2008) and Wilson et al. (2010), respectively. However, given the advective nature of an eastern boundary current and frequency of non-steady state conditions, it is difficult to generalize from our results to the rest of the Pacific. Clearly additional studies are necessary.

Since direct *in situ* measurements of subducted flux remains a technical challenge (Bishop et al., 2004; Omand et al., 2015), few models have attempted to explicitly include subducted POC into the carbon budget. Here we incorporated estimates of subducted flux for all 9 cycles based on results from a data-assimilative, Lagrangian particle study (Stukel et al., 2018c). By placing constraints on the subduction of POC to depth, we were better able to constrain the role of mesozooplankton in export production.

### 5. Conclusions

The LIEM used here incorporated numerous *in situ* measurements made during quasi-Lagrangian experiments in the CCE in order to constrain carbon flows through the ecosystem. These observations were made in water parcels spanning a wide range of conditions from highly productive upwelling regions to an oligotrophic offshore domain and consistently found that active transport of carbon by mesozooplankton was important to supplying the mesopelagic carbon demand. Our results suggest that passive POC flux—i.e. the portion of flux that is often assumed to dominate the biological pump—contributed 14% - 78% (median: 54%) of carbon flux. This finding has implications for the interpretation of sediment trap and ^234^Th disequilibrium measurements and for helping to reconcile a long-studied imbalance in the observed mesopelagic carbon budget. The LIEM also highlights the central importance of zooplankton in marine food webs and biogeochemistry. Excretion by vertical migrants is important in meeting bacterial carbon demand, while predation on vertical migrants supports mesopelagic resident fish communities. Our analysis comprises a unique, fully-resolved phytoplankton-to-fish coupled food web of the epipelagic and mesopelagic ocean. Nevertheless, substantial uncertainties remain, and targeted studies are necessary to validate the suggested relationships *in situ* and to test their applicability across the global ocean.

## Appendix A

**Table A1.** Measurement constraints used in the LIEM. Values given show the mean (µ) and 1 SD (σ) for each cycle except for min/max constraints which are blank. Marked values (*) were assumed values calculated from cycles of the same classification (see Section 2.1.2). All values are given in mg C m^−2^ d^−1^.

| Quantity |  | P07.1 |  | P0704-2 |  | P0704-4 |  | P0810-1 |  | P0810-2 |  | P0810-3 |  | P0810-4 |  | P0810-5 |  | P0810-6 |  |
| --- | --- | --- | --- | --- | --- | --- | --- | --- | --- | --- | --- | --- | --- | --- | --- | --- | --- | --- | --- |
| | | $\mu$ | $\sigma$ | $\mu$ | $\sigma$ | $\mu$ | $\sigma$ | $\mu$ | $\sigma$ | $\mu$ | $\sigma$ | $\mu$ | $\sigma$ | $\mu$ | $\sigma$ | $\mu$ | $\sigma$ | $\mu$ | $\sigma$ |
| 1 | C-14PP | 1233 | 74 | 587 | 35 | 2314 | 139 | 554 | 33 | 484 | 29 | 893 | 54 | 674 | 40 | 1672 | 100 | 325 | 19 |
| 2 | Delta PHY | -322 | 209 | 18 | 253 | -80 | 602 | 538 | 1680 | -1 | 50 | -76 | 166 | -399 | 4771 | -325 | 173 | -5 | 62 |
| 3 | Microzooplankton Grazing | 659 | 238 | 717 | 135 | 1282 | 225 | 472 | 301 | 243 | 51 | 357 | 74 | 277 | 41 | 498 | 48 | 138 | 43 |
| 4 | SMZ Grazing | 2249 | 1132 | 123 | 37 | 856 | 652 | 21 | 9 | 90 | 26 | 199 | 53 | 48* | 21* | 836* | 688* | 85* | 37* |
| 5 | LMZ Grazing | 93 | 87 | 42 | 18 | 123 | 55 | 18 | 12 | 14 | 9 | 66 | 26 | 21* | 15* | 51* | 55* | 19* | 14* |
| 6 | vmSMZ Grazing | 2092 | 907 | 137 | 44 | 669 | 499 | 50 | 30 | 110 | 34 | 249 | 113 | 72* | 50* | 753* | 538* | 101* | 47* |
| 7 | vmLMZ Grazing | 232 | 171 | 42 | 18 | 132 | 42 | 19 | 11 | 20 | 10 | 51 | 25 | 19* | 14* | 95* | 101* | 22* | 16* |
| 8 | SMZ + LMZ Grazing | 2342 | 1077 | 166 | 57 | 979 | 763 | 35 | 31 | 104 | 44 | 265 | 65 | 67* | 38* | 887* | 668* | 104* | 61* |
| 9 | Total MesoZ Grazing | 2306 | 1137 | 203 | 82 | 622 | 238 | 86 | 42 | 165 | 11 | 336 | 163 | 107* | 70* | 809* | 650* | 150* | 41* |
| 10 | Sed Trap @ 100m | 144 | 13 | 32 | 6 | 170 | 41 | 74 | 11 | 69 | 13 | 78 | 7 | 149 | 36 | 127 | 22 | 107 | 5 |
| 11 | Thorium @ 100m | 77 | 11 | 32 | 10 | 121 | 45 | 32 | 60 | 51 | 29 | 18 | 22 | 46 | 38 | 53 | 13 | 54 | 8 |
| 12 | Fecal Pellet Flux @ 100m | 135 |  | 4 |  | 54 |  | 4 |  | 8 |  | 5 |  | 35 |  | 14 |  | 4 |  |
| 13 | Minimum Subduction | 39 |  | 47 |  | 25 |  | 11 |  | 33 |  | 13 |  | 26 |  | 10 |  | 31 |  |
| 14 | Maximum Subduction | 79 |  | 55 |  | 45 |  | 33 |  | 47 |  | 15 |  | 51 |  | 19 |  | 59 |  |
| 15 | Epi Bacterial Prod (0-100m) | 66 | 27 | 22 | 12 | 53 | 30 | 240 | 137 | 80 | 17 | 148 | 21 | 351 | 60 | 400 | 37 | 101 | 8 |
| 16 | Min dBAC BP | 16 |  | 5 |  | 8 |  | 16 |  | 13 |  | 14 |  | 27 |  | 102 |  | 21 |  |
| 17 | Max dBAC BP | 52 |  | 16 |  | 27 |  | 52 |  | 41 |  | 45 |  | 89 |  | 336 |  | 69 |  |
| 18 | Deep NM Resp (dMYC) | 2.8 | 5.5 | 3.4 | 1.0 | 2.2 | 0.7 | 4.6 | 2.2 | 3.6 | 1.7 | 3.8 | 0.8 | 3.8 | 4.4 | 6.7 | 0.9 | 7.3 | 1.8 |
| 19 | Deep NM Poop (dMYC) | 1.2 | 2.4 | 1.5 | 0.5 | 1.0 | 0.3 | 2.1 | 1.0 | 1.6 | 0.7 | 1.7 | 0.4 | 1.7 | 1.9 | 3.0 | 0.4 | 3.3 | 0.8 |
| 20 | Deep NM Mort (dMYC) | 2.8 | 5.5 | 3.4 | 1.1 | 2.2 | 0.7 | 4.7 | 2.3 | 3.7 | 1.7 | 3.9 | 0.8 | 3.9 | 4.4 | 6.8 | 1.0 | 7.4 | 1.8 |
| 21 | Epi VM Resp (vmMYC) | 7.5 | 14.7 | 2.8 | 2.2 | 12.2 | 4.8 | 12.0 | 2.8 | 2.8 | 0.6 | 14.5 | 4.1 | 8.0 | 4.8 | 8.9 | 1.0 | 5.5 | 2.0 |
| 22 | Deep VM Resp (vmMYC) | 9.5 | 18.6 | 3.5 | 2.8 | 15.5 | 6.0 | 14.9 | 3.4 | 3.0 | 0.7 | 16.9 | 5.0 | 9.7 | 6.0 | 10.6 | 0.8 | 6.2 | 2.1 |
| 23 | Deep VM Poop (vmMYC) | 5.1 | 10.0 | 1.9 | 1.5 | 8.3 | 3.2 | 8.1 | 1.9 | 1.8 | 0.4 | 9.5 | 2.8 | 5.3 | 3.3 | 5.9 | 0.5 | 3.5 | 1.3 |
| 24 | Deep VM Mort (vmMYC) | 6.0 | 11.7 | 2.2 | 1.8 | 9.8 | 3.8 | 9.5 | 2.2 | 2.1 | 0.5 | 11.1 | 3.2 | 6.3 | 3.8 | 6.9 | 0.6 | 4.1 | 1.5 |

**Table A2.** Mesozooplankton biomass and minimum respiration estimates used in the LIEM. Respiration is given in mg C m^−2^ d^−1^ and biomass in mg C m^−2^.

| *Quantity* | *P0704-1* | *P0704-2* | *P0704-4* | *P0810-1* | *P0810-2* | *P0810-3* | *P0810-4* | *P0810-5* | *P0810-6* |
| --- | --- | --- | --- | --- | --- | --- | --- | --- | --- |
| Epipelagic Min Resp (SAR) | 8.0 | 8.6 | 8.0 | 0.0 | 0.0 | 0.0 | 0.0 | 8.9 | 0.0 |
| SMZ Min Resp | 24.4 | 5.1 | 20.6 | 4.8 | 26.3 | 18.0 | 36.3 | 31.6 | 27.3 |
| vmSMZ Min Resp | 5.3 | 0.4 | 5.5 | 16.4 | 3.3 | 8.6 | 0.0 | 21.5 | 0.0 |
| dSMZ Min Resp | 2.6 | 2.6 | 5.7 | 9.4 | 2.5 | 7.1 | 1.1 | 14.5 | 3.3 |
| LMZ Min Resp | 7.0 | 3.3 | 11.9 | 5.5 | 10.4 | 5.7 | 7.8 | 3.0 | 5.8 |
| vmLMZ Min Resp | 19.9 | 1.5 | 63.2 | 24.7 | 14.9 | 50.9 | 11.1 | 64.2 | 11.8 |
| dLMZ Min Resp | 3.2 | 5.4 | 6.7 | 20.4 | 7.6 | 13.3 | 4.9 | 31.3 | 13.5 |
| GEL Min Resp | 0.2 | 0.1 | 0.1 | 0.3 | 0.7 | 0.2 | 0.0 | 0.0 | 0.7 |
| dGEL Min Respiration | 0.4 | 0.1 | 0.5 | 0.4 | 1.1 | 0.3 | 0.1 | 0.4 | 0.8 |
| SMZ Biomass | 654 | 106 | 511 | 106 | 833 | 509 | 1249 | 1891 | 1346 |
| vmSMZ Biomass | 104 | 33 | 115 | 722 | 127 | 282 | 0 | 1120 | 0 |
| dSMZ Biomass | 499 | 280 | 715 | 968 | 307 | 834 | 111 | 1430 | 381 |
| LMZ Biomass | 424 | 173 | 478 | 971 | 586 | 1126 | 867 | 938 | 1763 |
| vmLMZ Biomass | 557 | 347 | 5175 | 993 | 1557 | 9859 | 1681 | 6122 | 1671 |
| dLMZ Biomass | 5928 | 2839 | 5702 | 7180 | 1818 | 3579 | 1395 | 7134 | 3942 |

## Acknowledgements

The authors would like to thank the crews and captains of the *R/V Thompson* and *Melville* for their superlative assistance in collecting the wide variety of observations used in this study. We thank our colleagues in the CCE for their continued support, energy and commitment to long-term ecological research of the pelagic. Additionally, the open source R community provided valuable technical support, and both directly and indirectly assisted in the development of the code used for the calculations presented here. This work was supported by NSF Biological Oceanography grants to the CCE LTER Program: OCE-0417616, OCE-1026607, OCE-1637632, and OCE-1614359. The source code and data used for the LIEM can be freely obtained under the MIT open-source license at https://github.com/tbrycekelly/Inverse_DVM or by contacting TBK.

## Author contributions

ML was responsible for cruise design and protistan zooplankton data. MO was responsible for mesozooplankton data. RG was responsible for phytoplankton data. PD was responsible for myctophid data. MS was responsible for particle export data. TK and MS designed the model. TK wrote the manuscript. All authors contributed to editing the manuscript.

