## Supplemental Information for "The Importance of Mesozooplankton Diel Vertical Migration for Sustaining a Mesopelagic Food Web"

**Supplemental Tables and Figures: The Importance of Mesozooplankton Diel Vertical Migration for Sustaining a Mesopelagic Food Web**

*Thomas B. Kelly, Peter C. Davison, Ralf Goericke, Michael R. Landry, Mark D. Ohman, Michael R. Stukel*

**Detailed Methods**

*LIEM Approach*

The MCMC approach begins with an initial solution that fits the equalities and inequalities (we started with the L_2_ solution, e.g. (Vézina and Piatt, 1988). A bounded random walk is then performed using the open source library *limSolve* (Soetaert et al., 2017). During each step of the random walk, a tentative solution is first produced. For this tentative solution to be accepted and for the random walk to continue from this new point, the inequality constraints need to be assessed to ensure all conditions are met. If the tentative solution lies outside the bound of an inequality constraint, the solution is reflected back into the valid solution space (i.e. mirror algorithm). The tentative solution is assessed using a weighted sum of squared residuals (SSR) between the model's solution and the *in situ* measured values including measurement uncertainty (Eq. 1).

$prob\left( x \right)=e^{-\frac{1}{2}\sigma^{-2}\left( Ax-b \right)^{T}\left( Ax-b \right)}$	(Eq 1)

where $\sigma$ is the measurement uncertainty and $Ax-b$ is the model-predicted *in situ* values.

A stochastic algorithm uses the ratio of the tentative solution’s probability (i.e. ${prob(x}_{n+1})$) and the previous solution's probability to determine if the tentative solution should be accepted (Eq. 2). If the solution is accepted, then the random walk procedure repeats from this new solution; but if it isn't, then the random walk procedure will start again from the previous solution.

$\frac{prob(x_{n+1})}{prob(x_{n})} \geq rand(0\to1)$	(Eq 2)

The random walk was performed for 200 million steps following a burn-in period of 20 million steps (code adapted from Van den Meersche et al., 2009). The burn-in period allowed the model to move away from the initial solution before sampling for the final solution set. Since the optimal acceptance ratio for high dimensional MCMCs has been reported to be around 25% (Roberts et al., 1997; Roberts and Rosenthal, 2001), the jump length for each cycle was adjusted to approximate that acceptance ratio for the sake of efficiency and consistency across cycles. The random walk solution is then subsampled yielding 10,000 solutions that fit the equality and inequality constraints while approximating the measurement constraints. The mean and 95% confidence intervals of these sets were used to determine the maximum likelihood solution and uncertainty for each output variable. When discussing ranges in flow values across multiple cycles, the values shown are the range in the mean solution for each cycle unless otherwise indicated.

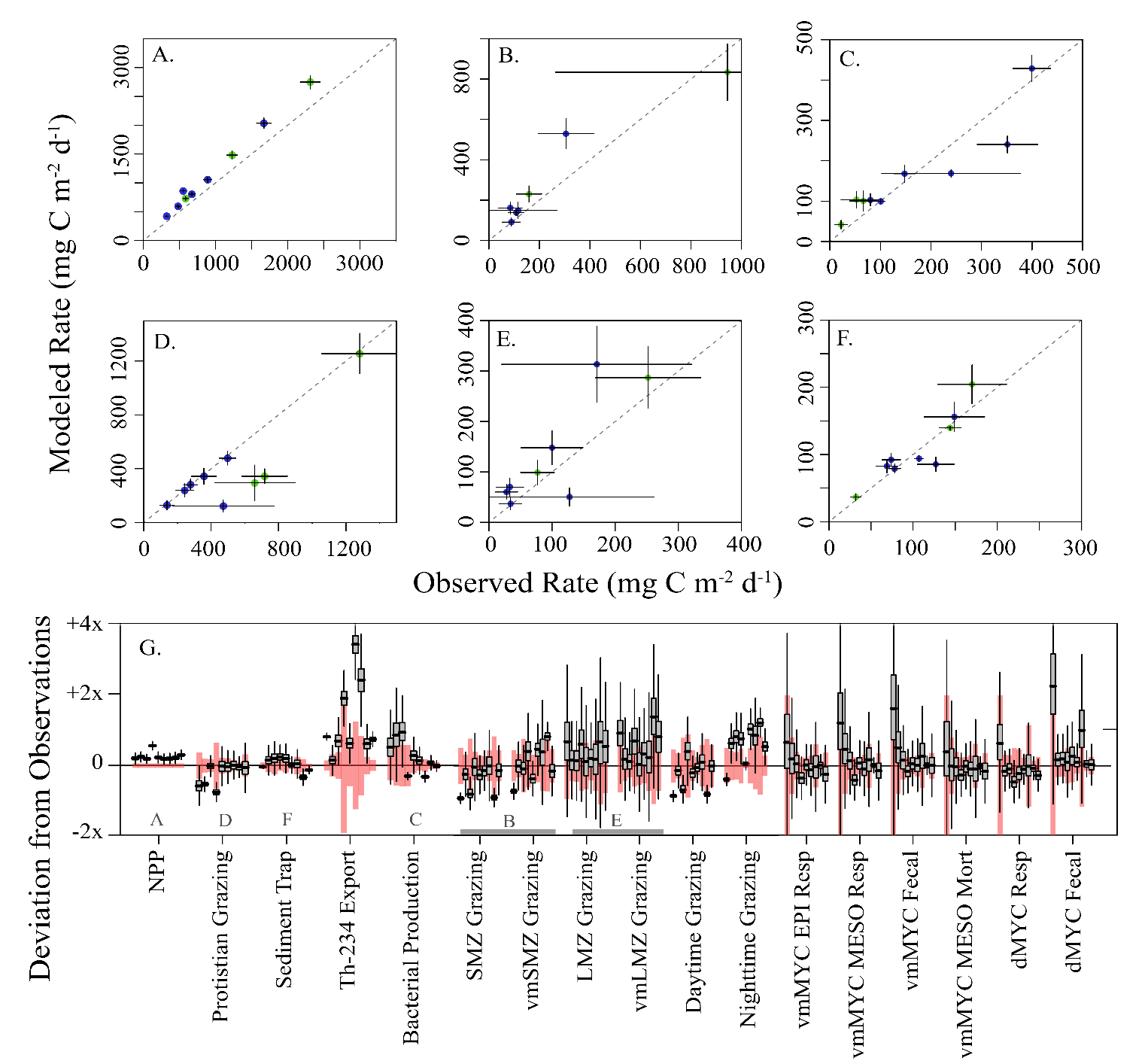

**Supplemental Figure 1.** Selected modeled rates verses observations: (A) primary productivity and export, (B) small mesozooplankton grazing, (C) epipelagic bacterial production, (D) protistian zooplankton grazing, (E) large mesozooplankton grazing, and (F) sediment trap export at 100m. Dashed line in each panel is a 1:1 reference line and error bars show 1 SD. (G) Box and whisker plot of relative model deviations from observations for each cycle (box shows inter-quartile range and whiskers to 95% CI). The cycles are shown in chronological order for each equation, and observations are as labeled. The red shading shows 1 SD of the observation (i.e. $\sigma_{obs}$).

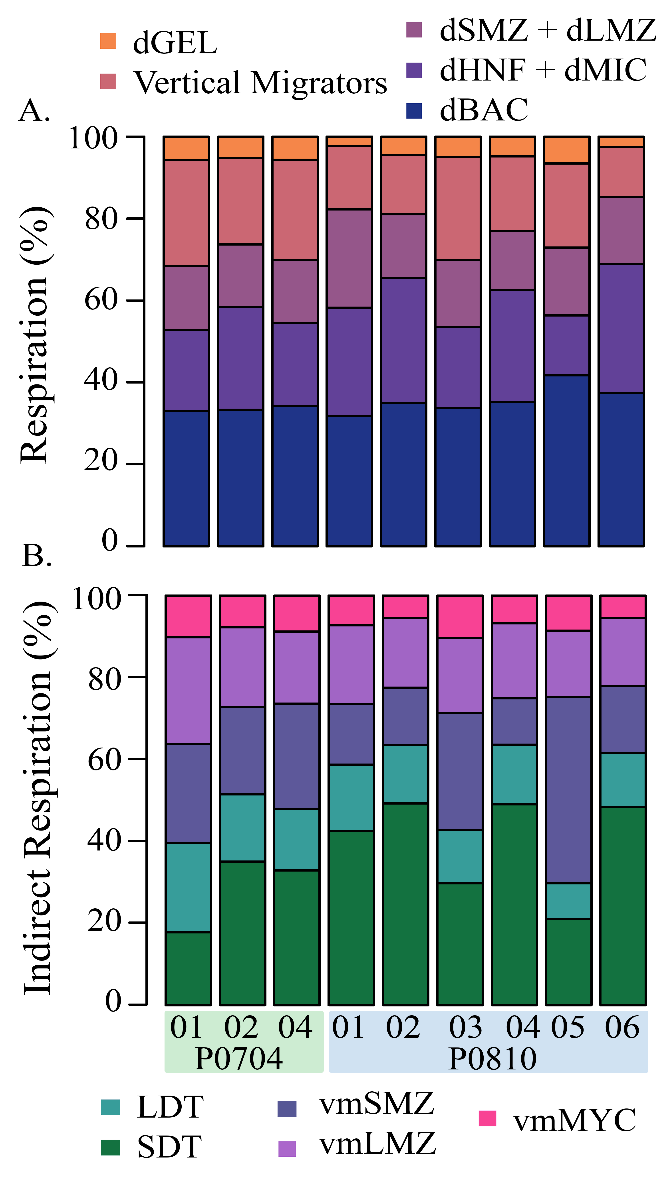

**Supplemental Figure 2.** (A) Proportion of mesopelagic respiration carried out by each category of mesopelagic organism. (B) Relative proportion of mesopelagic respiration supplied by each of the export pathways: passive transport by SDT and LDT, active transport by vmSMZ, vmLMZ, and vmMYC. Each were calculated from the indirect analysis.

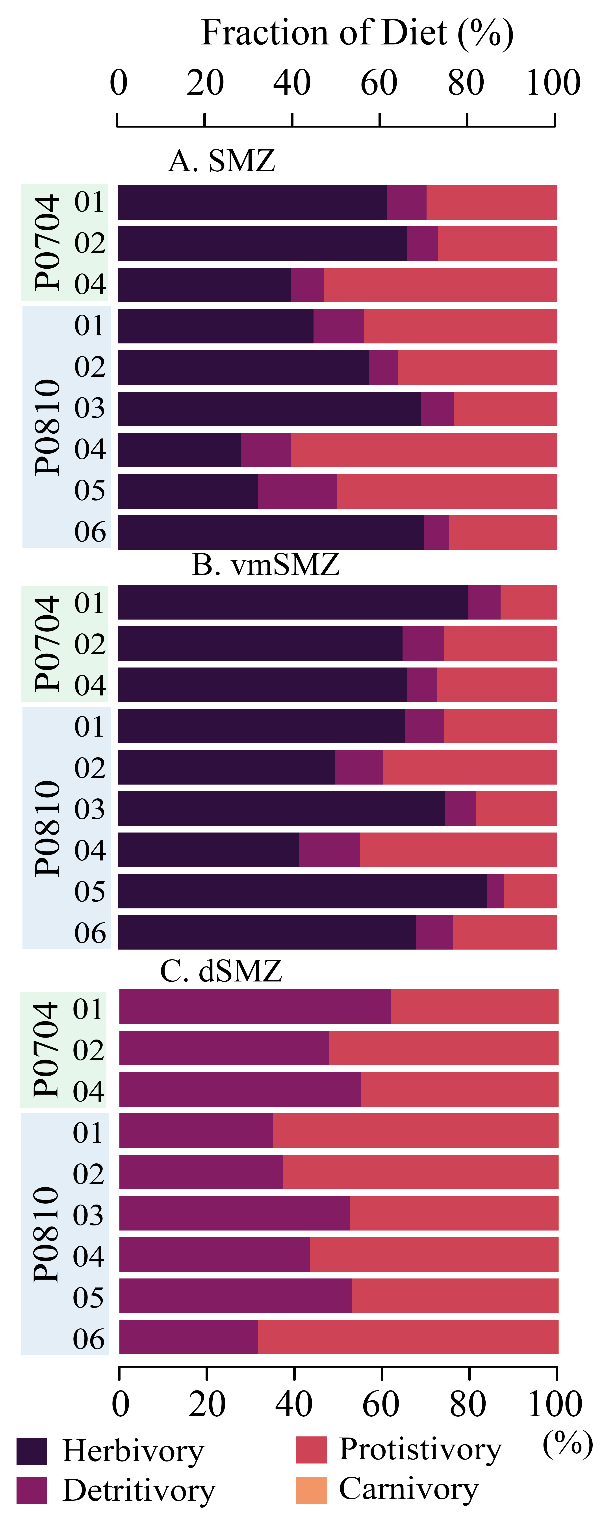

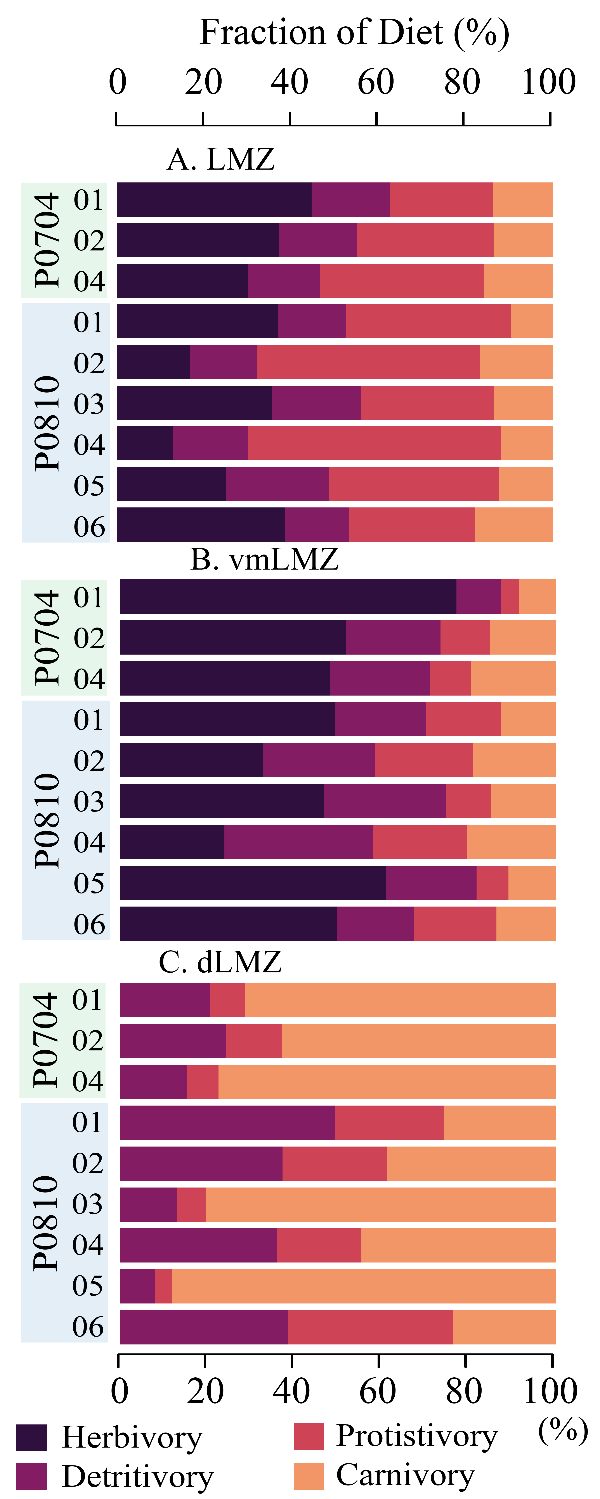

**Supplemental Figure 3**. The diets of each mesozooplankton compartment with small mesozooplankton on the left and large mesozooplankton on the right. (A) Resident epipelagic mesozooplankton, (B) diel vertically migrating mesozooplankton, and (C) resident mesopelagic mesozooplankton.

**Supplemental Table 1.** Input equations for the model and the associated organisms where G = grazing, R = respiration, D = detritus production, E = excretion of DOC, GPP = gross primary production, and NPP = net primary production. Subscript x signifies organism x, and epi and meso signify the epipelagic or mesopelagic flux, respectively. Measured constraints and mass balance constraints are provided in Appendix A. Activity specific constraints for vertically migrating mesozooplankton include epipelagic and mesopelagic temperatures (T­_epi_ and T_meso_, respectively) and a = 0.0648^o^C^-1^ (Ikeda, 1985).

|  | **#** | **Equation** | **Organism**  (where X= ) |
| --- | --- | --- | --- |
| GGE | 1a | $0.90\cdot G_{x}>R_{x}+D_{x}+E_{x}$ | MIC, HNF, dMIC, dHNF, SMZ, vmSMZ, dSMZ, LMZ, vmLMZ, dLMZ, Gel, dGEL |
|  | 1b | $0.60\cdot G_{x}<R_{x}+D_{x}+E_{x}$ |  |
| Bacterial Growth Efficiency | 2a | $0.95\cdot G_{x}>R_{x}+D_{x}+E_{x}$ | BAC, dBAC |
|  | 2b | $0.70\cdot G_{x}<R_{x}+D_{x}+E_{x}$ |  |
| Assimilation Efficiency | 3a | $0.50\cdot G_{x}>D_{x}$ | HNF, dHNF, MIC, dMIC, SMZ, vmSMZ^†^, dSMZ, LMZ, vmLMZ^†^, dLMZ, GEL, dGEL |
|  | 3b | $0.10\cdot G_{x}<D_{x}$ |  |
| Excretion | 4a | $0.10\cdot G_{x}<E_{x}$ | HNF, dHNF, MIC, dMIC, SMZ, vmSMZ, dSMZ, LMZ, vmLMZ, dLMZ, GEL, dGEL |
|  | 4b | $R_{x}<E_{x}$ |  |
|  | 4c | $\frac{E_{x,epi}}{E_{x,meso}}>\frac{\exp\left( a\cdot T_{epi} \right)}{exp(a*T_{meso})}$ | vmSMZ, vmLMZ |
| Phytoplankton Excretion | 5a | $E_{x}>0.02 NPP$ | PHY |
|  | 5b | $E_{x}<0.55 NPP$ |  |
| Respiration from Biomass | 6a | $R_{x}>in situ estimate$ | SMZ, vmSMZ, dSMZ, LMZ, vmLMZ, dLMZ, GEL, dGEL |
|  | 6b | $\frac{R_{x,epi}}{R_{x,meso}}>\frac{\exp\left( a\cdot T_{epi} \right)}{exp(a*T_{meso})}$ | vmSMZ, vmLMZ |
| Respiration from Ingestion | 7 | $R_{x}>0.20\cdot I_{x}$ | HNF, dHNF, MIC, dMIC, SMZ, vmSMZ, dSMZ, LMZ, vmLMZ, dLMZ, GEL, dGEL |
| Phytoplankton Respiration | 8a | $R_{x}>0.10\cdot GPP$ | PHY |
|  | 8b | $R_{x}<0.55\cdot GPP$ |  |
| Phytoplankton Excretion + Resp | 9a | $E_{x}+R_{x}>0.29 GPP$ | PHY |
|  | 9b | $E_{x}+R_{x}<0.62 GPP$ |  |
| Deep Bacterial Production | 10a | $I_{x}-R_{x}>BP minimum$ | dBAC |
|  | 10b | $I_{x}-R_{x}<BP maximum$ |  |
| Fecal Pellet Flux | 11 | (LDT to dLDT) > minimum | LDT, dLDT |
| Vertical Migrations | 12 | (GEL to dGEL) > minimum | GEL, dGEL |
| Subduction | 13a | (subducted detritus) > minimum | SDT, dSDT, LDT, dLDT |
|  | 13b | (subducted detritus) < maximum |  |

**Supplemental Table 2.** Listing of flows and mean flux values from each cycle (mg C m^-2^ d^-1^).

Supplemental_table2.xlsx

**Supplemental Appendix SA**

Spreadsheets used in the model and mirrored at <http://github.com/tbrycekelly/Inverse-DVM>.

1. Model.xlsx
2. Constraints.xlsx
